# Determining off-target effects of splice-switching antisense oligonucleotides using short read RNAseq in neuronally differentiated human induced pluripotent stem cells

**DOI:** 10.1101/2025.01.09.631923

**Authors:** EC Kuijper, L van der Graaf, BA Pepers, M Guimarães Ramos, S Korhorn, LJA Toonen, D Cats, RAM Buijsen, E Mina, H Mei, WMC van Roon-Mom

## Abstract

Antisense oligonucleotides (AONs) are small pieces of chemically modified DNA or RNA that bind to RNA in a sequence-specific manner based on Watson-Crick base-pairing. Splice-switching AONs are designed to modulate pre-mRNA splicing, thereby for instance restoring protein expression or modifying the eventual protein to restore its function or reduce its toxicity. Given the current lack of *in silico* methods that adequately predict off-target splicing events, assessment of off-target effects of AONs in human cells using RNAseq could be a promising approach. The identification and prioritization of potential off-target effects for validation and further investigation into the biological relevance would contribute to the development of safe and effective AONs. In this study, we used three different splice-switching AONs targeting three different human transcripts to study their transcriptome-wide, hybridization-dependent off-target effects with short read RNAseq. Using the computational tools rMATS and Whippet, we identified differential splicing events of which only a minority could be explained by hybridization, illustrating the difficulty of predicting off-target effects based on sequence homology. The main splicing events could all be validated with RT-PCR. Furthermore, from the three AONs studied, one AON induced considerably more changes in gene expression and splicing compared to the two other AONs assessed, which was confirmed in a validation experiment. Our study demonstrates that the interpretation of short read RNAseq data to determine off-target effects is challenging. Nonetheless, we showed that valuable results can be obtained as it allows the comparison of toxicity between different AONs within an experiment and identification of AON-specific off-target profiles.

## Introduction

Antisense oligonucleotides (AONs) are small pieces of chemically modified DNA or RNA that bind to RNA in a sequence-specific manner based on Watson-Crick base-pairing. AON therapies therefore have exceptional specificity and provide the possibility to target any transcript in the cell. This makes AON therapies particularly suitable for treating genetic diseases. At the moment, 16 AON therapies have been approved by the U.S. Food and Drug Administration and/or the European Medicines Agency (1). Different chemical modifications of the AON exist to improve affinity, stability, uptake and bioavailability of the AONs (2). In general, the phosphorothioate (PS) backbone modification is used to improve AON stability and uptake as it promotes binding to serum proteins with low affinity, preventing renal clearance. This however also drives toxicity as coagulation is inhibited and the complement system is activated. Modifications increasing the affinity to target RNA allow a lower dose, thereby reducing some of the PS-induced toxicity. Such modifications are modified sugar ribose (e.g. 2’O-methoxyethyl (MOE) and 2’O-methyl (2’OMe)) or completely modified nucleotides such as phosphorodiamidate morpholino oligomers (PMO). These modifications render the AON resistant to nucleases such as RNase H (2, 3). Depending on the chemical modality, AONs can act through three main mechanisms: protein knockdown, protein restoration and protein modification (2). For protein knockdown, gapmer AONs are most often used. Gapmer AONs are designed to induce RNase H-mediated degradation of the RNA and therefore contain a central ‘gap’ with unmodified DNA nucleotides to allow RNase H activity. Splice-switching AONs are completely modified and hence resistant to RNase H-mediated degradation. This allows splice-switching AONs to modulate pre-mRNA splicing, resulting in the inclusion or exclusion of an exon or use of an alternative splice site. This way, splice-switching AONs can restore protein expression or modify the resulting protein to restore its function or reduce its toxicity (2).

As with each type of drug, AONs can have off-target effects. These unintended events can be related to the AON chemistry (class effects) or the AON sequence and are divided into effects independent or dependent of AON hybridization to RNA (4). Hybridization-independent effects are immunostimulatory effects, thrombocytopenia, complement activation and inhibition of coagulation (4). The sequence of the AON can also affect these hybridization-independent effects as has been observed for immunostimulatory effects, such as CpG motifs activating TLR9, and thrombocytopenia (4). Hybridization-dependent off-target effects are the result of unintended complementary binding of the AON to RNA that share sequence homology with the intended target sequence. Theoretically, this type of off-target effect could be predicted based on *in silico* assessment of sequence homology (5, 6). Although BLAST is widely used to study sequence homology, it is not always reliable for short sequences (7, 8). It has been shown that by using GGGenome, a search engine for nucleotide sequences, to determine sequence homology or by predicting AON/RNA minimum free energy the prediction of potential off-targets can be improved (6, 8–10). As AONs can bind both RNA, pre-mRNA and non-coding RNA, these should all be included in the *in silico* assessment of sequence homology.

Various studies have evaluated *in silico* predicted hybridization-dependent off-target events *in vitro* in human cells. Here, we focus on splice-switching AONs. In a retrospective study of 105 RNA sequencing (RNAseq) experiments, the number of splicing changes observed in an exon decreased with an increasing number of mismatches between the exon and the AON (11). In line with the requirement for splice-switching AONs to bind at spliceosome-interfering sites, there was only a 16% probability of inducing a splicing event at perfectly complementary binding sites. Nonetheless, even when including predicted binding sites with up to 5 mismatches with the AON, only 40.6% of all splicing events could be explained, showing that there are more differential splicing effects than predicted by using sequence homology alone. Interestingly, the differential gene expression changes did not seem to be related to the number of mismatches, gaps or insertions, suggesting that off-target hybridization of the splice-switching AONs to the RNA was not the main cause for the differential gene expression (11).

Given the current lack of *in silico* methods that adequately predict off-target splicing events, assessment of off-target effects of AONs in human cells using RNAseq is a promising approach (11). For the detection of alternative splicing events in RNAseq data, a wide range of computational tools have been developed in recent years. Although quantitative analysis methods are being developed for long-read sequencing analysis, they are still more suitable for isoform discovery rather than of isoform quantification (12–14). Hence, in the current study we focused on splicing analysis of short-read sequencing data.

In general, three main types of tools can be distinguished. Isoform-based tools assess the level of individual transcript isoforms assembled with the use of isoform-specific reads. For short read sequencing technology, the low proportion of isoform-specific reads makes these tools less reliable (15–17). Secondly, exon-based tools use exon counts without their transcriptomic context. Although this decreases the complexity and required computational resources, the accuracy is reduced and the type of splicing event cannot be inferred (16–18). Finally, event-based methods are in general thought to be the most reliable for differential splicing analysis. Here, inclusion levels of predefined event patterns are assessed based on the ratio of reads that support inclusion or exclusion.

However, a standard approach for differential splicing analysis is still lacking and the sparsely described settings when using these different tools make comparison and reproduction of studies difficult. Given the low performance for all tools, it is usually recommended to use the overlap of different tools to eliminate tool-specific false positives and increase the validity of downstream analysis (15, 16). Recent studies comparing different splicing tools showed that rMATS was superior in detecting novel events and that the combination of rMATS with Whippet improved the false discovery rate (15, 16, 19).

In this study, we used three different splice-switching AONs targeting three different human transcripts to study their transcriptome-wide, hybridization-dependent off-target effects. The three splice-switching AONs were previously shown to be effective in cell and/or animal models and targeted either *APP*, *ATXN3* or *HTT* (Fig 1A). The *APP* gene is involved in Alzheimer disease and cerebral amyloid angiopathy and the APP-AON was designed to skip exon 17 (all exon numbers mentioned refer to the MANE select transcript) as this exon contains a mutation in Dutch-type CAA (20). *ATXN3* encodes the ataxin-3 protein, which contains a polyglutamine domain that is expanded in spinocerebellar ataxia type 3 (SCA3). The ATXN3-AON induces a skip of exon 10, which encodes the polyglutamine domain and leads to the formation of a truncated protein (21). The HTT-AON induces a partial exon skip of exon 12, which is identified as a use of an alternative 5’ splice site (22, 23). The eventual modification of the HTT protein is hypothesized to reduce the toxicity of mutant HTT involved in Huntington Disease (HD). As all three AONs are designed to target the brain, we used neuronal cells derived from control induced pluripotent stem cells (iPSCs) to mimic more closely the target tissue gene expression. We used rMATS in combination with Whippet for differential splice analysis to describe the extent of unintended splicing events after transfection of the splice-switching AONs, focusing on hybridization-dependent events, and evaluate this method for improving *in vitro* off-target identification pipelines.

**Figure 1.**
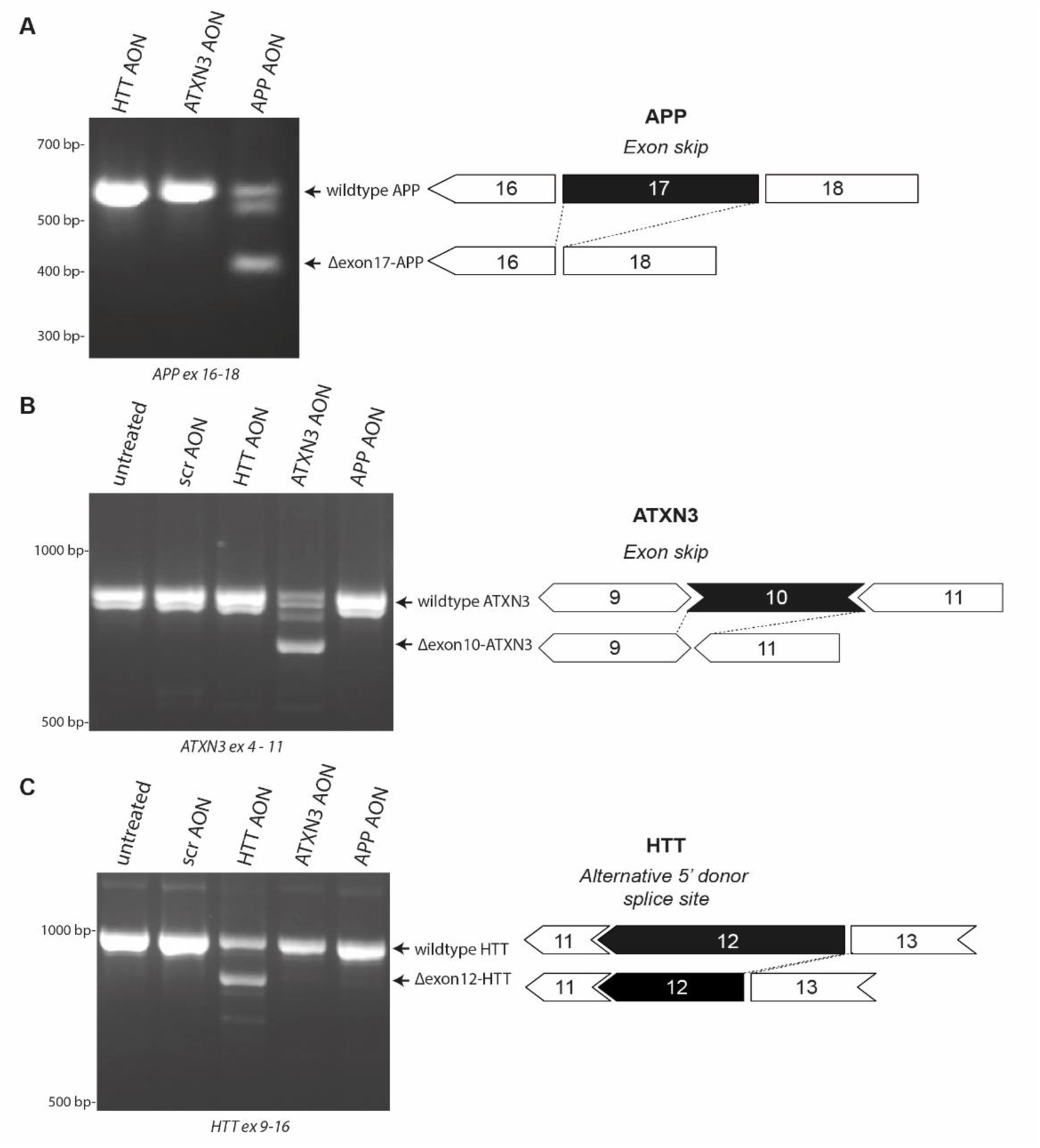
Intended events of three splice-switching AONs. The AONs included in this study were designed to skip exon 17 in APP (A) or exon 10 in ATXN3 (B) or induce the use of an alternative 5’ donor splice site in exon 12 of HTT (C). Using RT-PCR, the intended events in the target genes were confirmed upon transfection of the targeting AONs and were not observed in untreated cells or cells transfected with a scrambled (scr) AON or another targeting AON.

## Methods

### Antisense oligonucleotides

Antisense oligonucleotides (Table 1) were designed according to the guidelines published by Aartsma-Rus (24). Basic Tm was calculated by OligoCalc (25). Binding energy within the AON and between AONs was calculated by the Fold and bifold web servers from RNAstructure (https://rna.urmc.rochester.edu/RNAstructureWeb/index.html).

**Table 1.**
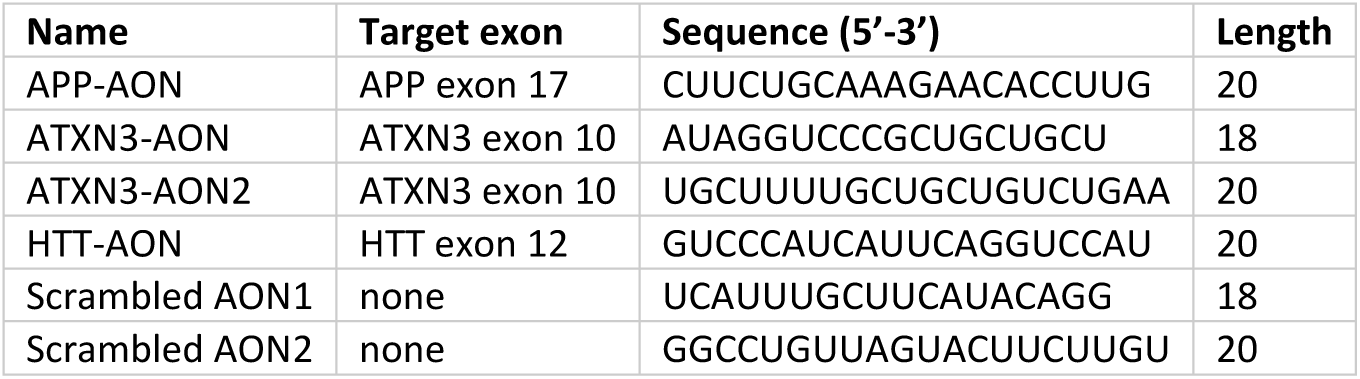
Antisense oligonucleotides used in this study.

### Neuronal culturing and transfections

Female healthy control human induced pluripotent stem cell lines C1.1 (LUMCi003-A), C1.2 (LUMCi003-B) and C2 (LUMCi023-A) were used to generate neuronal progenitor cells (NPCs) following previously described methods (26, 27). Cells were cultured at 37°C and 5% CO_2_. Differentiation of NPCs into neuronal cultures was done by plating 5 × 10^5^ NPCs in a PDL/laminin-coated 6-well plate in STEMdiff neuron differentiation medium (STEMCELL Technologies) with daily medium changes. After 1 week, 5 to 10 × 10^5^ cells were replated in PDL/laminin-coated 6-well plates in STEMdiff neuron differentiation medium, followed by half medium changes using BrainPhys medium every 2 to 3 days for neuronal maturation. At day 11 after the start of neuronal maturation, cells were transfected during 4 hours with AONs diluted to 200nM concentration in Opti-MEM medium (Thermo Fisher, Waltham, MA, USA) with 0.3% Lipofectamine 2000 (Life Technologies, Carlsbad, CA, USA). After this, medium was replaced to BrainPhys medium and cells were harvested 3 days later using ReleSR (STEMCELL Technologies) and spun down. Cell pellets were snapfrozen and stored at –20 °C until RNA isolation. For each transfection condition, 6 or 7 (C1.1), 3 (C1.2) or 4 (C2) replicates (wells) were included. The validation of potential off-target effects was performed in cell line C1.1 with 2 replicates.

### RNA isolation and library preparation

RNA was isolated from cell pellets using the Reliaprep RNA isolation kit (Promega, Madison, WI, USA) following manufacturer’s instructions. DNAse treatment was performed to degrade genomic DNA. RNA was sequenced in two batches, one containing the samples from cell line C1.1 and the other containing cell lines C1.2 and C2. Library preparation and RNA sequencing was performed at GenomeScan B.V. (Leiden, The Netherlands). The quality of RNA was assessed with the Fragment Analyzer (Agilent Technologies). Approximately 250 ng of total RNA was used as starting material, and the average RQN values were 9.9 (SD ± 0.3). To process the samples, the NEBNext Ultra Directional RNA Library Prep Kit was used for C1.1 and the NEBNext Ultra II directional RNA (including dual index/UMI) for C1.2 and C2. Sample preparation was performed according to the protocol “NEBNext Ultra Directional RNA Library Prep Kit for Illumina” (NEB #E7420S/L). Briefly, mRNA was isolated from total RNA using the oligo-dT magnetic beads. After fragmentation of the mRNA, a cDNA synthesis was performed. This was used for ligation with the sequencing adapters and PCR amplification of the resulting product. Quality of sequencing libraries was determined with the Fragment Analyzer to assess insert size and quantity. Pooled samples were clustered and sequenced using a Novaseq6000 (read lengths 2 × 150 cycles). Primary processing and base calling was performed with Illumina’s NCS1.4 and RTA3 for C1.1 and NCS1.8 and RTA3.4.4 for C1.2 and C2. Demultiplexing and generation of FASTQ files was done with Illumina scripts (bcl2fastq v2.2 for C1.1 and bclconvert v3.10.5 for C1.2 and C2). The FASTQ files will be uploaded to the European Genome-phenome Archive (EGA).

### Sequencing data processing

Processing of sequencing data was performed using the BIOPET Gentrap in-house pipeline (http://biopet-docs.readthedocs.io/en/v0.7.0/pipelines/gentrap/). The fastqc toolkit (v0.11.2) was used to evaluate sequencing quality (http://www.bioinformatics.babraham.ac.uk/projects/fastqc/). Sickle (v1.33 with default settings) and Cutadapt (v1.10) were used to clean up reads. Cleaned reads were aligned to the reference genome build 10 (GRCh38) using STAR aligner version 2.3.0e (28). The non-default settings used by STAR are “—outFilterMultimapNmax 1 –outFilterMismatchNmax 10 – outSJfilterReads Unique”. The gene raw read counts were generated through HTSeq (v0.6.1). Data quality control was performed by examining the per sequence GC content, GC coverage bias, adapter content, insert size and gene coverage. No samples were excluded on the basis of these metrics.

### Differential gene expression analysis

Differential gene expression analysis was performed in R (v4.0.2) using DESeq2 (v1.30.1). Because of the difference in sequencing depth and sample size between the two data sets, different filtering settings were applied. We selected genes with >0.15 fragments per million for at least 6 samples for dataset 1 from cell line C1.1. Data set 2 (C1.2 and C2) was split into two data sets, one for each cell line. Genes were filtered for >1 fragments per million for at least 3 (cell line C1.2) or 4 (cell line C2) samples. Differential expression was performed using the DESeq-function with default settings and results were extracted with disabled independent filtering. Genes with an adjusted P-value (Benjamini-Hochberg) of <0.05 were considered significantly differentially expressed. Official gene symbols and gene biotypes were assigned to ENSEMBL identifiers using the getBM function of biomaRt package (v 2.46.3). Pathway enrichment analysis was performed with gprofiler2 (v.0.2.1) using the fdr correction method and a significance level of 0.05.

### Differential splicing analysis

Differential splicing analysis was performed with rMATS and Whippet. The settings were adapted to allow the detection of novel splice sites and optimized as described in Supplementary File 1 and final settings are described here. Using rMATS turbo (v4.1.2), we performed comparisons between samples treated with scrambled AON (--b1) and samples treated with a targeting AON (--b2). The following settings were used: –-readLength 150, –-variable-read-length, –-cstat 0.05, –-AnchorLength 15, –-allow-clipping, –-novelSS. For cell line C1.1, –-libType was fr-firststrand and for cell lines C1.2 and C2, –-libType was fr-secondstrand. As input gtf-file, Homo_sapiens.GRCh38.109.gtf was used, but filtered for the genes used as input for differential gene expression analysis of cell line C1.1 (25124 genes in total) and filtered for isoforms annotated as gencode-basic. For further analysis, results based on junction counts were filtered in R, keeping only the events for which each sample had at least 10 informative reads (IJC + SJC). Events with FDR < 0.05 were considered as significant.

For splicing analysis using Whippet, merged BAM-files were made for all comparisons (scrambled AON vs. targeting AON) and used as input for the index together with the filtered gtf-file described above. For generating the index, bam-both-novel was disabled and bam-min-reads was 6. Quantification of each sample was performed using the default settings.

Next, we determined the overlap of significant events obtained by rMATS and Whippet for the event types identified by both tools: exon skipping (SE/CE), alternative splice donor site (A5’SS/AD), alternative splice acceptor site (A3’SS/AA) and retained intron (RI)(15, 16). To compare the output from rMATS and Whippet, a standardized genomic region was extracted for all splicing events as described by Olofsson et al (16). The overlap between the tools was determined and filtered using the scripts available within the *rmappet* package (https://github.com/didrikolofsson/rmappet/)(16). For each event, the average dPSI was calculated from the dPSI’s obtained by both tools. In the overlap, some of the events detected by Whippet were connected to multiple different events detected by rMATS, as only rMATS takes into account different flanking regions. To determine the overlap of events between conditions, we used the dplyr (1.0.8) functions to identify the events for which rMATS coordinates, rMATS flanking regions and Whippet coordinates were the same.

### GGGenome sequence homology

Predicted binding sites in the human prespliced RNA with a maximum distance of 3 were determined using GGGenome (https://gggenome.dbcls.jp/). The AON target sequence (reverse complement of AON sequence) was used as input to search into the human pre-spliced RNA database (RefSeq curated on GRCh38/hg38.p14, D3G Feb, 2023). A maximum number of 3 mismatches was allowed and only the plus strand was searched for. Next, we assessed the overlap between the events detected and the predicted binding sites by comparing the genomic locations. Using R, it was assessed whether the start or end coordinate of the predicted binding site was within the surrounding region of a splicing event, as given by the rMATS results. The region between the furthest ends of the flanking exons plus 200 bp were taken in to account, assuming that AON binding in this region most likely could affect splicing of the exon. For A5’SS and A3’SS events, only one flanking exon was available within the rMATS results for A5’SS and A3’SS events. Hence, we manually assessed the overlap of the predicted binding sites with the flanking region on the other side of these events.

### cDNA synthesis and PCR

The intended splicing events in *APP*, *ATXN3* and *HTT* as well as potential off-target splicing events were assessed with RT-PCR. To that end, cDNA was synthesized from RNA with Transcriptor First Strand cDNA synthesis kit (Roche) using random hexamer primers. Subsequently, the PCR was run with the primers listed in Table 2 and the following program: initiation 4 minutes 95°C; 36 cycles of 30 seconds 95°C denaturation, 30 seconds 59°C annealing, 1 minute 72°C extension; 7 minutes 72°C final elongation. PCR products were run on an agarose gel and visualized with the OptiGo-750 (Isogen Life Science). The intensity of the bands was quantified in ImageJ and the skip percentage was calculated by dividing the intensity of the skipped band by the intensity of all bands.

**Table 2.**
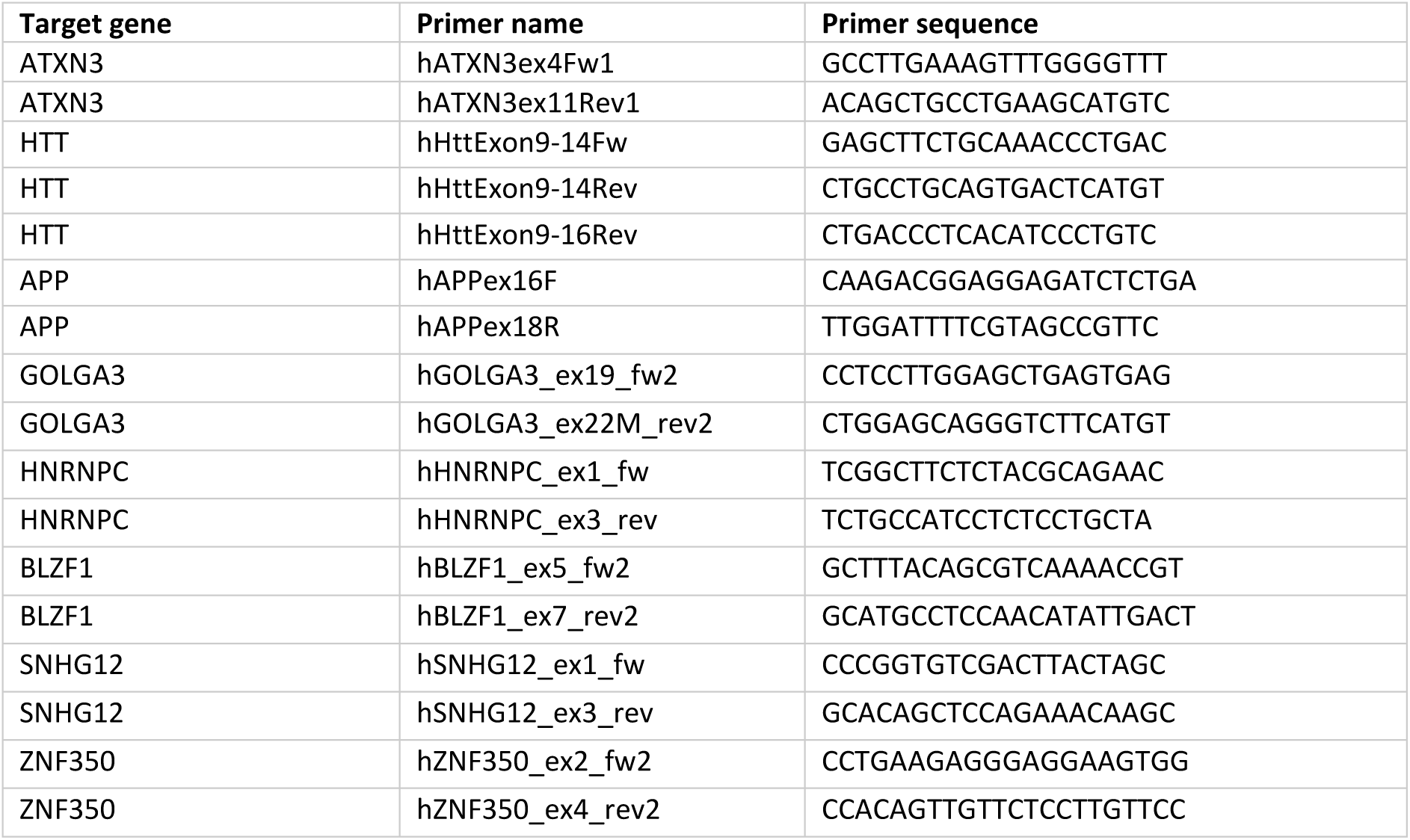
List of primers used in this study.

### Visualization and statistics

Graphs were made in R using ggplot2 (v.3.3.6) and statistics were applied using ggpubr (v.0.4.0). Venn diagrams were made in R with VennDiagram (v.1.7.3).

## Results

### Intended AON-induced splicing events were confirmed in iPSC-derived neuronal cells

iPSC-derived neuronal cells from control cell line C1.1 were transfected with a scrambled AON or an AON designed to induce exon skipping within APP, ATXN3 or HTT (Fig 1). Cell death was observed in the ATXN3-AON-transfected samples as was confirmed by a lower RNA yield (Fig S1A). To confirm successful transfection, the intended exon skip events in APP, ATXN3 or HTT were verified using RT-PCR, which showed about 40% of skip efficiency for all AONs, as was expected based on previous studies (Fig 1)(20–23).

### AON-transfection led to massive gene expression changes

To first explore the transcriptome-wide effects after AON transfection, we performed differential gene expression analysis. After RNA sequencing, the percentage of reads assigned to a gene was significantly lower for the ATXN3-AON-transfected samples compared to the other conditions whereas the percentage of reads that aligned to a region without any features (“no feature”) was significantly higher (Fig S1B-D). Nonetheless, the mean number of assigned reads did not differ between the groups and the average number of assigned reads among all samples was about 43 million (Fig S1E). After filtering out the genes with a low expression level, we tested 25,124 genes in total. To exclude AON class effects, we compared each targeting AON with the scrambled AON. All three AONs caused both significant up– and downregulation of genes (Table 3, Fig 2A, Supplementary File 2). The APP-AON showed the lowest number of differentially expressed genes (DEGs): 3911. The HTT-AON depicted 11,387 DEGs and most DEGs were observed with ATXN3-AON: 16,453. Especially regarding DEGs with an absolute log fold change >1, the ATXN3-AON caused major gene deregulation compared to the APP-AON and HTT-AON. Mainly protein coding genes were changed, but also lncRNAs and miRNAs were affected, particularly by the ATXN3-AON and HTT-AON (Fig 2B).

**Figure 2.**
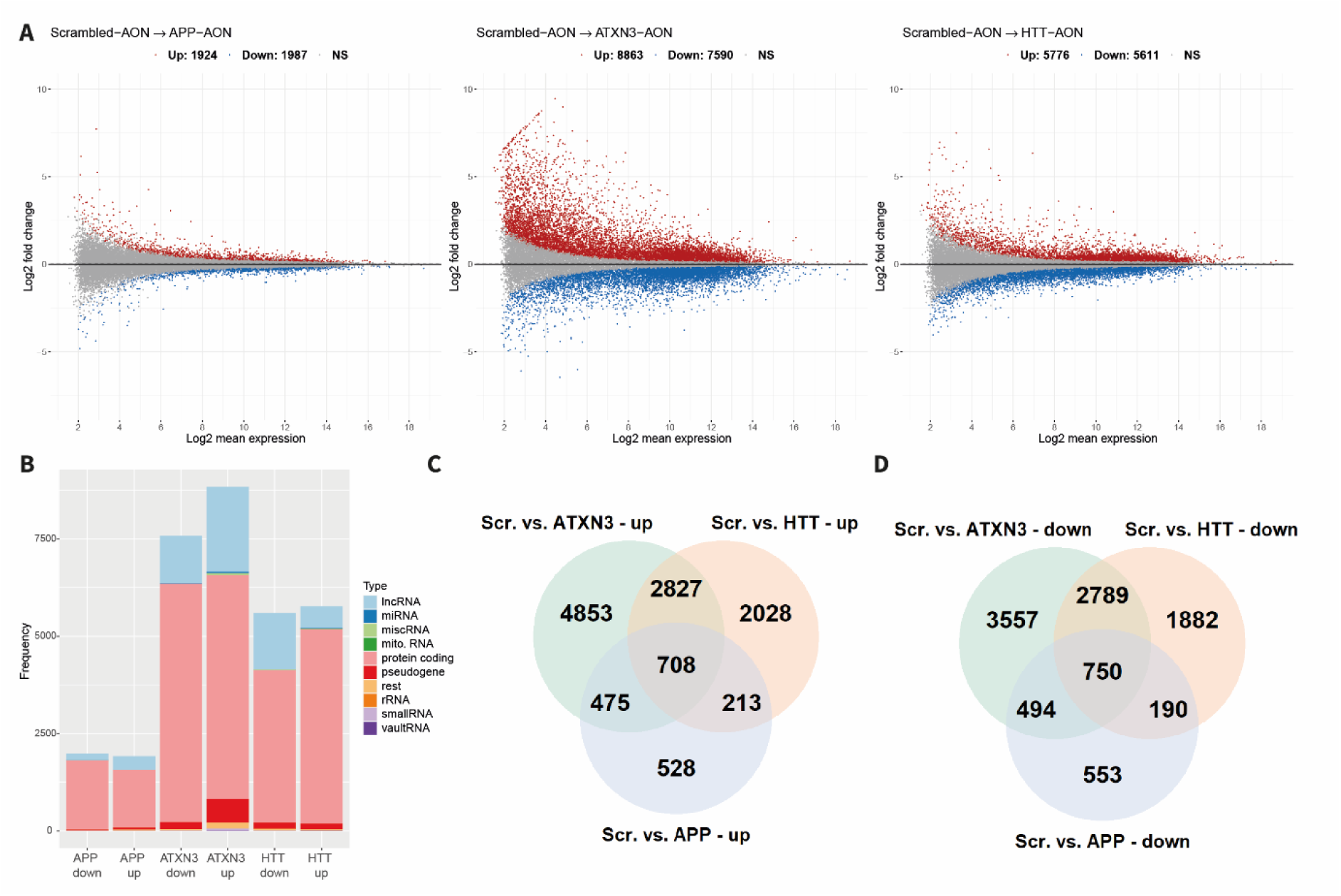
Differential gene expression analysis showed massive gene expression changes upon AON transfection. A) Fold change and gene expression level of significantly upregulated (red) and downregulated (blue) genes and non significant genes (grey) for each targeting AON compared to transfection with the scrambled AON, showing largest gene deregulation upon transfection with ATXN3-AON. B) Number of significantly down-or upregulated genes upon transfection with each AON per gene biotype demonstrated that mainly protein coding RNA and lncRNA were differentially expressed. C-D) Number of overlapping up– and downregulated genes for the three targeting AONs.

**Table 3:** Number of genes differentially expressed between cells transfected with scrambled AON or one of the targeting AONs (APP, HTT, ATXN3). LFC: log fold change.

| Comparison | Significant | Up | Up, \|LFC\|>1 | Down | Down, \|LFC\|>1 |
| --- | --- | --- | --- | --- | --- |
| Scrambled AON vs. APP-AON | 3911 | 1924 | 206 | 1987 | 94 |
| Scrambled AON vs. HTT-AON | 11387 | 5776 | 778 | 5611 | 841 |
| Scrambled AON vs. ATXN3-AON | 16453 | 8863 | 3857 | 7590 | 1664 |

To gain insight in the effects of AON transfection, we performed pathway enrichment analysis. First, we assessed the genes that were deregulated by all three AONs, which were 708 upregulated and 750 downregulated genes (Fig 2C-D). Pathway analysis for the overlapping upregulated genes demonstrated enrichment of for instance cellular biosynthetic processes, regulation of RNA metabolism and regulation of transcription (Fig S2, Supplementary File 3). Pathways that were enriched among downregulated genes were related to cell signaling and neuronal projection development (Fig S2).

Next, we studied the AON-specific effects, which might be related to side effects or downstream effects of the intended events (Fig S3, Supplementary File 3). Downregulated genes by the APP-AON were for instance related to apoptosis, cell cycle, cellular organization and biosynthetic processes. There were no significantly enriched pathways among the upregulated genes for the APP-AON. Genes upregulated by the HTT-AON showed enriched pathways such as cell cycle and cell division, whereas pathways enriched among the downregulated genes were related to RNA metabolism and regulation of transcription. An opposite pattern was observed by genes deregulated by ATXN3-AON, as upregulated genes were related to RNA metabolism and regulation of transcription, while downregulated genes were related to cell cycle and cell division.

### Splicing events identified by rMATS and Whippet were highly correlated

To study the transcriptome-wide effects on splicing, we performed differential splicing analysis using rMATS and Whippet comparing each targeting AON with the scrambled AON. The output of both tools is given as percent spliced in (PSI), indicating the percentage of reads that contain the indicated exon. When comparing the different AON conditions, the difference in PSI is indicated as deltaPSI (dPSI), which is equivalent to the exon skip percentage. Since the dPSI from both tools showed a correlation of 0.93 (APP-AON), 0.87 (ATXN3-AON) and 0.90 (HTT-AON), indicating that results were consistent and comparable, the average deltaPSI from both tools was used for further assessments (Fig 3A). In total, when compared to the scrambled AON, 489 events were found by both tools for the APP-AON, 6230 events for the ATXN3-AON and 867 events for the HTT-AON, distributed over all chromosomes (Table 4, Fig 3B, Supplementary File 4). Exon skipping events were most abundant and also showed the highest overlap between the tools in terms of percentage (Supplementary File 1).

**Figure 3.**
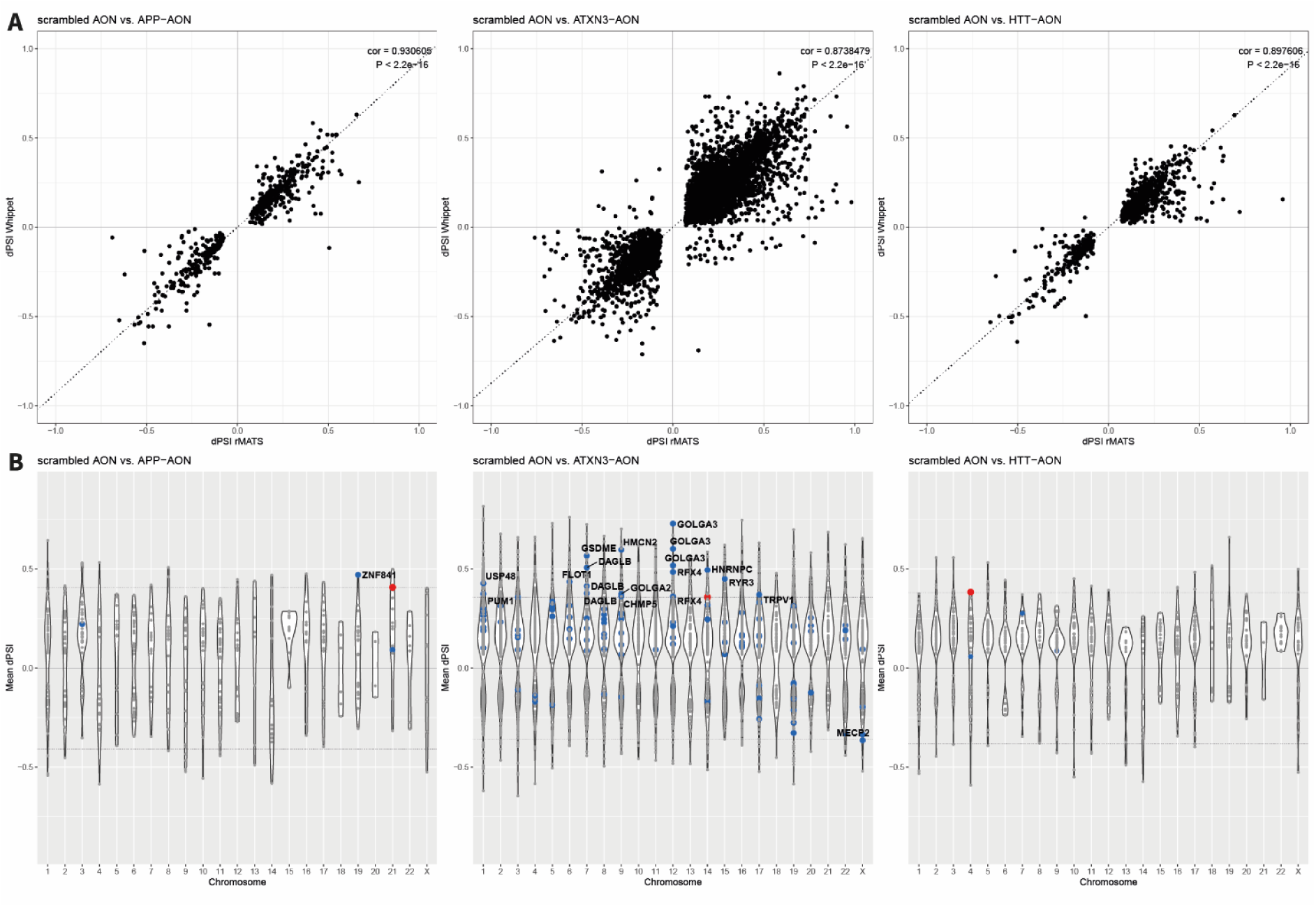
Differential splicing events found by both rMATS and Whippet. A) The delta percent spliced in (dPSI) identified by rMATS and Whippet for a specific splicing event showed a strong correlation. B) Splicing events were distributed over all chromosomes. The red dot indicates the intended splicing event for the targeting AON in *APP*, *ATXN3* and *HTT* respectively. Potentially hybridization-dependent events are indicated by blue dots and labelled when the absolute mean dPSI was larger than the intended event.

**Table 4.** Number of significant results obtained by both rMATS and Whippet per event type.

| Event type | APP-AON | ATXN3-AON | HTT-AON |
| --- | --- | --- | --- |
| Exon skipping | 468 | 5741 | 842 |
| Alternative splice donor site | 9 | 286 | 14 |
| Alternative splice acceptor site | 10 | 142 | 9 |
| Retained intron | 2 | 61 | 2 |
| *Total* | *489* | *6230* | *867* |

### Intended splicing events were found by both rMATS and Whippet

First, we assessed the intended exon skip events of the AONs in APP, ATXN3 and HTT as a positive control. For all AONs the intended exon skip event was the only significant event found in the respective target gene (Table S1, Supplementary File 4). The dPSI levels were 0.36 for ATXN3-AON, 0.38 for HTT-AON and 0.41 for APP-AON (Fig 3B, red dots) and corresponded well with the observed skipping efficiency of about 40% as detected with RT-PCR. All events were highly significant as indicated by the FDR of 0 by rMATS and a probability of >0.97 by Whippet. The scrambled AON treated samples demonstrated no natural occurrence of the intended splicing events, showing a PSI of about 1.

### Hybridization-dependent off-target splicing events do not play a major role after transfection

Next, we aimed to identify potentially hybridization-dependent splicing events by selecting the splicing events that 1) could be linked to a predicted binding site of the AON in the transcriptome and 2) were AON-specific. To that end, we first determined predicted binding sites in the human prespliced RNA with a maximum number of 3 mismatches with the AON using GGGenome (Table 5, Supplemental File 5). As expected, for the targeting AONs the only binding sites found with 0 or 1 mismatches (including gaps or insertions) were the intended binding sites. Compared to APP-AON and HTT-AON, ATXN3-AON showed more predicted binding sites, which might be related to the shorter size of this AON as it contains 18 nucleotides whereas APP-AON and HTT-AON both consist of 20 nucleotides (6). The scrambled AON, with a length of 18 nucleotides as well, showed the highest number of predicted binding sites including 2 binding sites with 1 mismatch. These 2 predicted binding sites were located in *DCT* and *IL1RAPL1*, but no differential splicing events were detected in these genes in all three comparisons.

**Table 5:** Predicted binding sites found for AONs of interest. The number of predicted binding sites containing 0, 1, 2 or 3 mismatches (including gaps or insertions) are listed as well as the total number of predicted binding sites and the number of genes these binding sites contained. * = intended binding site.

| AON | 0 mismatches | 1 mismatch | 2 mismatches | 3 mismatches | Total binding sites | Total genes involved |
| --- | --- | --- | --- | --- | --- | --- |
| Scrambled AON | 0 | 2 | 160 | 5353 | 5515 | 3480 |
| APP-AON | 1* | 0 | 26 | 613 | 640 | 556 |
| ATXN3-AON | 0 | 1* | 78 | 1782 | 1861 | 1470 |
| HTT-AON | 1* | 0 | 13 | 324 | 338 | 313 |

To assess the link between predicted binding sites and the observed splicing events, we analyzed whether the predicted binding sites occurred in the regions surrounding the significant splicing events that we observed (Table 6). Only a low proportion of the splicing events could be linked to a predicting binding site, suggesting that many splicing events are not induced in a hybridization-dependent manner. Including the target site, less than 2% of total events could be related to predicted binding sites of the APP-AON (5 out of 489, 1.0%), ATXN3-AON (122 out of 6230, 1.9%) and HTT-AON (5 out of 867, 0.58%). Subsequently, we assessed the second criterium that hybridization-dependent events should be AON-specific. Interestingly, nearly all of the significant splicing events that could be linked to a predicted binding site of the targeting AON were specific for the AON involved. We identified 4 potentially hybridization-dependent events for the APP-AON, 117 for the ATXN3-AON and 4 for the HTT-AON (Fig 3B, blue dots, Table 6). Furthermore, it is important to note that there were also events matching to predicted binding sites of the scrambled AON, including some AON-specific events (Table 6). Since the scrambled AON was present in all comparisons, events based on hybridization of the scrambled AON are expected to arise in all three conditions. However, power differences because of different group sizes or indirect effects of the targeting AON may complicate data comparisons and interpretation.

**Table 6.**
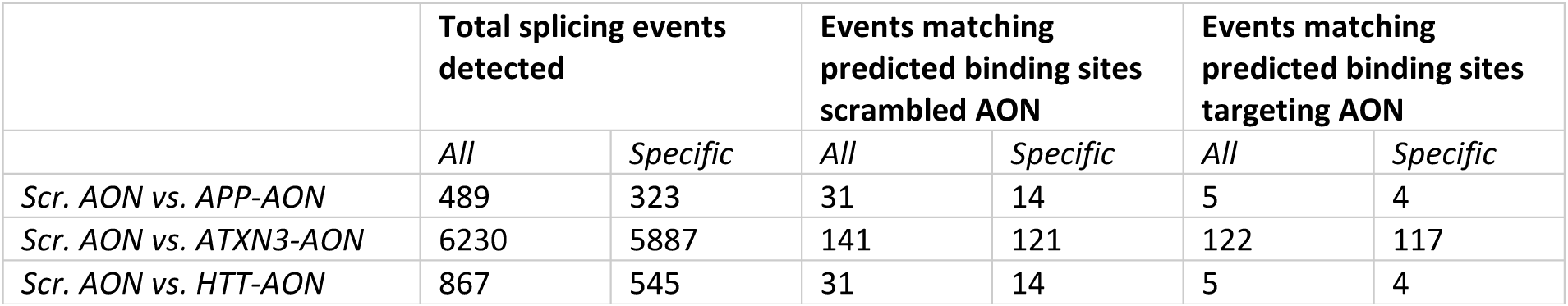
Only few of the splicing events detected could be linked to a predicted binding site based on AON sequence homology. Both the number of all splicing events detected and the number of AON-specific splicing events are shown.

We further studied the potentially hybridization-dependent events of the targeting AONs and identified the events that were larger than the intended event of the AON. We validated the largest event in an independent transfection experiment in C1.1 using RT-PCR. The potentially hybridization-dependent events for APP-AON (apart from the intended event) were located in *ZNF841*, a transcription factor, *QTRT2*, which is involved in tRNA modification, and *TTC3*, an E3 ubiquitin ligase. The splicing event located in *ZNF841* showed a larger dPSI of 0.47 compared to the intended splicing event of the AON. For this event, interference of the AON with splicing would be likely as the predicted binding site of the AON is on the border of exon 2, the exon that was skipped. The AON-specific skip was confirmed using RT-PCR, although several bands were visible that could not be explained (Fig 4A). In the differential gene expression analysis, we found that *ZNF841* expression levels were significantly downregulated (LogFoldChange=-0.71, FDR= 1.30E-11). As the exon skipped within the transcription factor *ZNF841* is part of the 5’UTR, it might be involved in regulating its expression levels.

**Figure 4.**
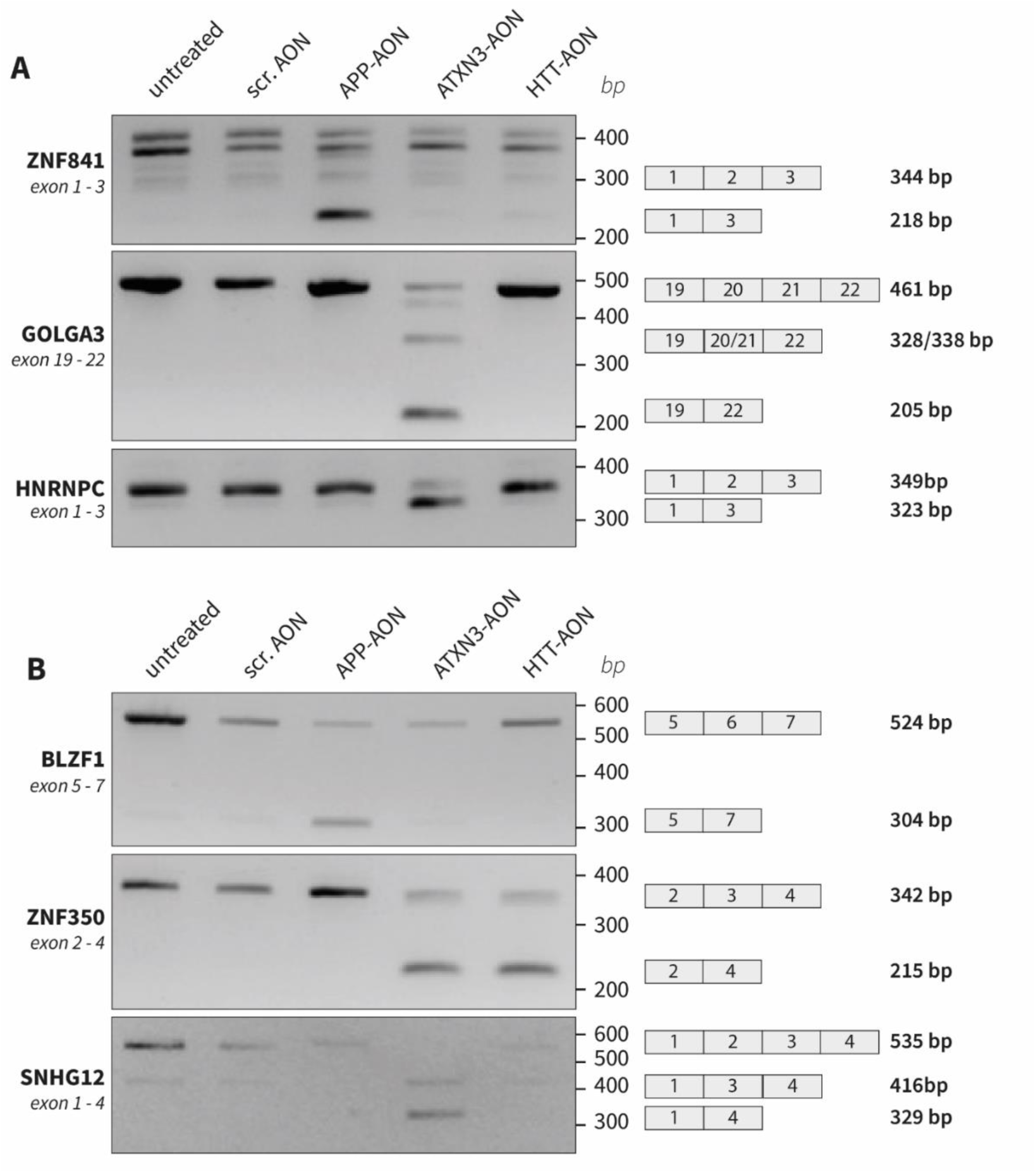
Validation of splicing events of interest. Using RT-PCR, the largest hybridization-dependent events (A) and overall largest splicing events (B) detected were confirmed.

For HTT-AON, apart from the event in *HTT*, the potentially hybridization-dependent events were located in *MAN1B1*, an enzyme involved in N-glycan trimming in the ER-associated degradation pathway, *CDK13*, involved in the cell cycle but also associated with mRNA processing, and *WDR1*, involved in disassembly of actin filaments. These splicing events all had a smaller dPSI than the intended splicing event in *HTT*.

For ATXN3-AON, the genes showing potentially hybridization-dependent events were mainly related to cellular component organization, neurogenesis and RNA processing, and included the splice regulating genes *HNRNPA1*, *HNRNPC*, *SF3A3* and *SUPT6H*. From the 116 potentially hybridization-dependent events other than the intended event, 19 events located in 14 genes showed a larger dPSI than the intended event (Table S2). The two events with the largest dPSI were found in *GOLGA3*, involving the skip of either exon 21 (dPSI = 0.73) or exon 20 (dPSI = 0.60) between flanking exons 19 and 22, which together could reflect the skipping of both exon 20 and 21. Furthermore, another large potentially hybridization-dependent splicing event was found in *GOLGA3*, involving the skip of exon 21 between exon 20 and 21 (dPSI = 0.52). The predicted binding site of the AON was located in exon 21. RT-PCR showed a product in which both exon 20 and 21 were skipped, as well as a product that reflects the individual skip of exon 20 (133bp) or exon 21 (123bp), which sizes are too similar to give a conclusive answer (Fig 4A). Among the 19 larger events was also the skip of exon 2 (dPSI = 0.49) in *HNRNPC*, a gene involved in splicing, that was interesting as it was linked to predicted binding sites of both the ATXN3-AON and the scrambled AON. The predicted binding site of ATXN3-AON was located in the middle of the intron between exon 2 and 3, where the AON might interfere with recognition of the branch point. The scrambled AON was predicted to at a distance of more than 500 bp from exon 1, where interference with splicing is less likely but not ruled out. Moreover, this event was interesting as a transcript lacking exon 2 is expressed naturally. Validation using RT-PCR nicely showed the natural occurrence of a transcript lacking exon 2 as well as the increase upon ATXN3-AON transfection specifically (Fig 4A), supporting the link to the predicted binding site of ATXN3-AON.

In conclusion, only a small proportion of the splicing events detected are potentially hybridization-dependent off-target events. In addition, these in general showed a smaller effect size than the intended event, which might be related to the fact that all predicted binding sites had at least 2, but mostly 3 mismatches with the AON sequence.

### Majority of major splicing events were not hybridization-dependent

Apart from several potentially hybridization-dependent events described above, other events showing a higher absolute dPSI than the intended splicing event were observed as well (Fig 3B). Given their biological relevance as unexpected but large effects on splicing, we subsequently looked into these events as well and aimed to validate the largest events with RT-PCR in a separate experiment. For the APP-AON, 34 splicing events were found in this category of which 20 were specific for the APP-AON. Two high frequency APP-AON events in *LPAR1* matched with a predicted target of the scrambled AON but were only found upon APP-AON transfection. The event with the largest dPSI was found in *BLZF1*, encoding for Basic Leucine Zipper Nuclear Factor 1, showing a skip of exon 6 by the APP-AON that was not found for the other 2 AONs, as was confirmed with RT-PCR (Fig 4B).

The HTT-AON induced 41 larger events of which 19 were AON-specific and none matched with predicted targets of the scrambled AON (Fig 3B). The event with the largest absolute dPSI (dPSI = 0.66) was the skip of exon 3 in *ZNF350*, an event that also occurred upon ATXN3-AON transfection where it showed a dPSI of 0.70. Both were observed with RT-PCR (Fig 4B).

In the cells treated with ATXN3-AON, from the 715 larger events, 654 events were unique for transfection with the ATXN3-AON (Fig 3B). Moreover, 11 events showed overlap with a predicted binding site of the scrambled AON. The event with the largest dPSI (0.82) was found in *SNHG12*, a long non-coding RNA, and involved the skip of exon 3 of the MANE transcript. However, the flanking regions indicated by rMATS suggested that it concerned skipping of this exon within the T218 transcript (ENST00000692541.1), which lacks exon 2. An RT-PCR for exon 1 to 4 indeed showed that after ATXN3-AON transfection, a lower band appeared reflecting the skip of both exon 2 and 3 confirming the splicing event detected (Fig 4B).

Interestingly, especially for APP-AON and HTT-AON, there was a large overlap among the events showing a larger dPSI than the intended event of the AON as 12 events were present in all three conditions. Large events shared by all three conditions were located in *ARHGAP29*, *CDC16*, *CNN3, IFTAP*, *INTS6L*, *LGR4, PIKFYVE*, *PRXL2C*, *RAPGEF2*, *VTI1B*, *YME1L1* and *YY1AP1* and all showed a negative dPSI that was similar for the three AONs (Table S3).

### Validation experiment confirmed ATXN3-AON-related transcriptomic changes

From the three AONs investigated, the ATXN3-AON showed cell death upon transfection, lower numbers of mapped reads and widespread transcriptomic changes in terms of both differential splicing and differential gene expression. Furthermore, the numbers of potentially hybridization-dependent events and splicing events that showed a larger dPSI than the intended event were much higher than for the APP-AON and HTT-AON. Pathway analysis demonstrated that pathways related to cell cycle and cell division were downregulated. To validate these signs of toxicity for ATXN3-AON, we performed a follow-up experiment in a different clone from cell line C1 (C1.2) as well as in an additional cell line C2 using a different scrambled AON (scrambled AON2). In addition, we included a newly designed AON that induced the same exon 10 skip in *ATXN3* but with a different sequence, to evaluate whether the widespread effects of ATXN3-AON were sequence-specific and not related to the skipping of exon 10. When we compared the predicted binding sites of ATXN3-AON with ATXN3-AON2, there were less predicted binding sites for ATXN3-AON2 (Table 7, Supplementary File 5).

**Table 7.** Predicted binding sites found for the AON sequences used in C1.2 and C2. The number of predicted binding sites containing 0, 1, 2 or 3 mismatches (including gaps or insertions) are listed as well as the total number of predicted binding sites and the number of genes these binding sites contained. * = intended binding site.

| AON | 0 mismatches | 1 mismatch | 2 mismatches | 3 mismatches | Total binding sites | Total genes involved |
| --- | --- | --- | --- | --- | --- | --- |
| Scrambled AON2 | 0 | 0 | 6 | 234 | 240 | 219 |
| ATXN3-AON | 0 | 1* | 78 | 1782 | 1861 | 1470 |
| ATXN3-AON2 | 1* | 2 | 41 | 987 | 1031 | 885 |

Similar to what was observed in cell line C1.1, cell death upon transfection and a lower number of reads assigned to a gene were observed for ATXN3-AON in cell line C2, but less so in cell line C1.2. (Fig S4, Fig S5). For ATXN3-AON2, no cell death was observed. There were on average 16.0M reads in C1.2 and 9.91M reads in C2 (Fig S5F). Given the differences in gene expression between C1.2 and C2 as observed with principal component analysis, we performed the differential gene expression analysis for the cell lines separately (Fig S6). The exon skip in *ATXN3* was confirmed by RT-PCR analysis for both ATXN3-AONs in C1.2 and C2 (Fig S7). Due to the difference in sequencing depth between the experiment in cell line C1.1 and this validation experiment in C1.2 and C2, only the Whippet tool could significantly identify the intended exon 10 skip event for ATXN3-AON (dPSI=0.26 in line C1.2 and dPSI=0.23 in line C2) and ATXN3-AON2 (dPSI=0.11 in both lines).

Next, we focused on validating the differential gene expression results (Table 8, Supplementary File 6). When we looked at the results in the three cell lines for ATXN3-AON, in total there were 119 upregulated DEGs and 49 downregulated DEGs shared by all three cell lines (Fig 5A-B), supporting that these are related to ATXN3-AON transfection. The 49 downregulated DEGs showed enrichment of GO biological processes related to for instance lipid metabolism and cell differentiation, which were identified as well in cell line C1.1 (Supplementary File 7). Then we compared the DEGs after transfection with ATXN3-AON and ATXN3-AON2. There were less DEGs for ATXN3-AON2, with 9 upregulated genes and 42 downregulated in both cell lines C1.2 and C2 (Table 8). Only 3 upregulated genes (*DBNDD1*, *NCS1* and lncRNA ENSG00000260804) and 6 downregulated genes (*FDFT1*, *IRAG1*, *HMGCS1*, *MYLK* and two lncRNAs ENSG00000275620 and ENSG00000231107) were shared between ATXN3-AON and ATXN3-AON2 (Fig 5C-D). Except for IRAG1, the identical genes were dysregulated in cell line C1.1 by APP-AON and/or HTT-AON as well and hence not specifically related to the exon 10 skip in *ATXN3*.

**Figure 5.**
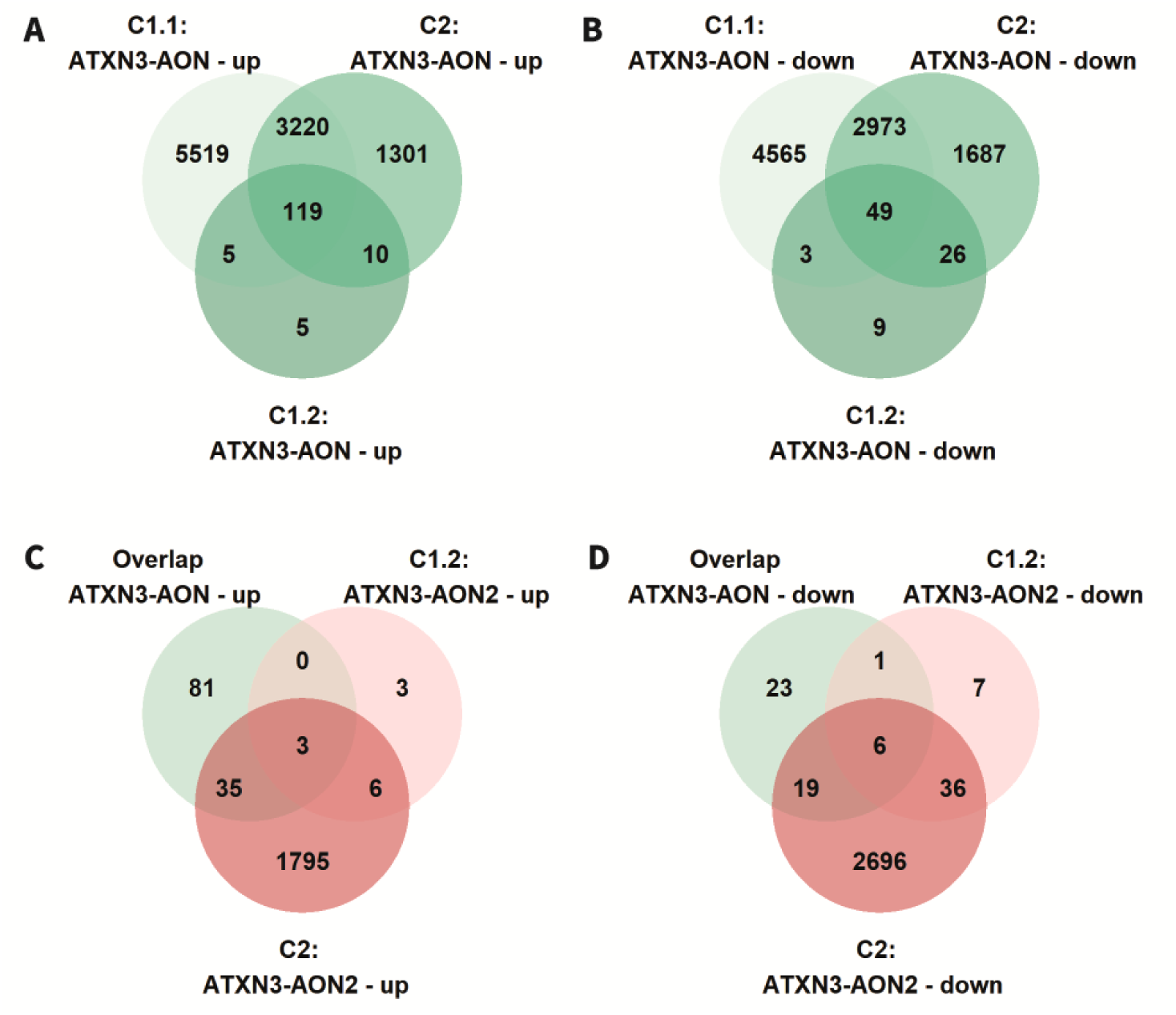
Shared differentially expressed genes upon ATXN3-AON transfection. The overlap of differentially expressed genes for ATXN3-AON in cell lines C1.1, C1.2 and C2 showed 119 shared upregulated genes (A) and 49 shared downregulated genes (B). C-D) When the shared deregulated genes for ATXN3-AON were compared to the genes differentially expressed upon ATXN3-AON2 in cell lines C1.2 and C2, only a small overlap was observed.

**Table 8.**
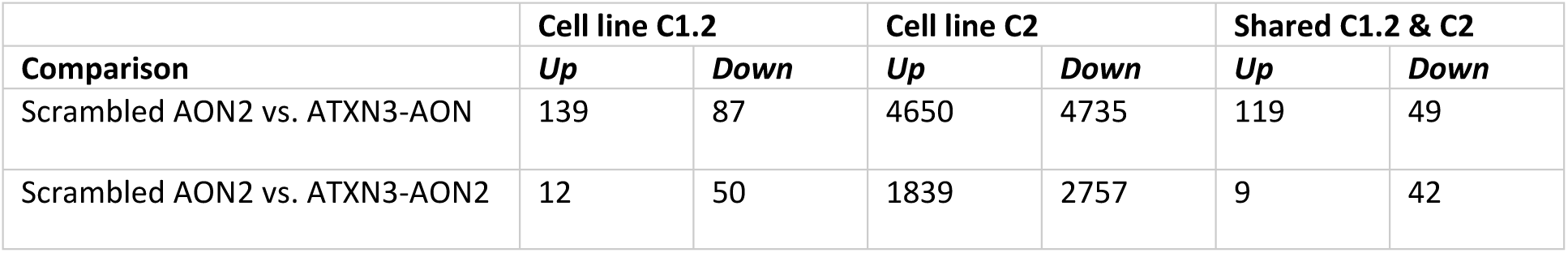
Number of genes differentially expressed between transfection with scrambled AON2 versus transfection with ATXN3-AON10.4, ATXN3-AON10.a or ATXN3-AON10.h in cell line C1.2 and cell line C2.

After analyzing the DEGs, we assessed the overlap of all splicing events induced by ATXN3-AON in the different cell lines (Table 9, Supplementary File 8). Out of the 6230 splicing effects found in the first experiment with cell line C1.1, 198 were detected in all 3 cell lines (Fig 6A). These 198 shared events occurred in 178 genes that showed overrepresentation of genes involved in RNA splicing and neuron differentiation upon pathway analysis (Supplementary File 9). The dPSI of all shared events showed a strong correlation between the cell lines (Fig S8). From the 198 shared events, 96 events (48%) were identified in C1.1 as AON-specific events larger than the intended event in *ATXN3*. Similar to the differential gene expression analysis described above, although the number of events was the lowest in C1.2, the majority of the events that we could detect in C1.2 was shared with the other cell lines (Fig 6A). This showed that although sequencing depth and sample size influence the number of overlapping splicing events, the main AON-induced splicing events could be replicated in the three cell lines.

**Figure 6.**
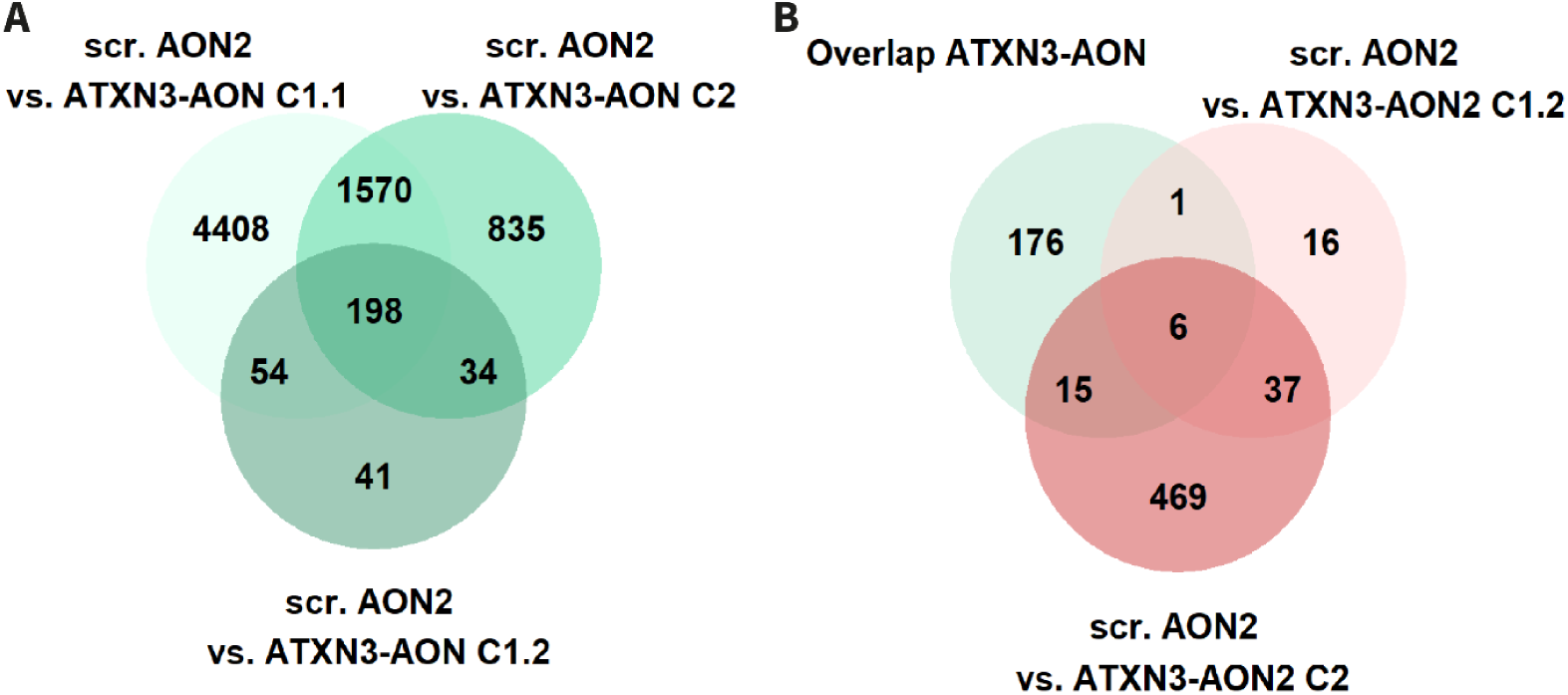
Shared splicing events upon ATXN3-AON transfection. A) ATXN3-AON showed 198 splicing events shared among cell lines C1.1, C1.2 and C2. B) From these 198, 6 were induced by ATXN3-AON2 as well in both cell lines C1.2 and C2.

**Table 9:** Significant splicing events for ATXN3-AON and ATXN3-AON2 in cell lines C1.2 and C2 compared to scrambled AON2. Results from ATXN3-AON transfection in cell line C1.1 from experiment 1 are shown for comparison.

|  | ATXN3-AON |  |  | ATXN3-AON2 |  |
| --- | --- | --- | --- | --- | --- |
| Cell line: | C1.2 | C2 | C1.1 | C1.2 | C2 |
| Exon skipping | 316 | 2462 | 5741 | 59 | 478 |
| Alternative splice donor site | 9 | 86 | 286 | 1 | 18 |
| Alternative splice acceptor site | 1 | 65 | 142 | 0 | 6 |
| Retained intron | 1 | 24 | 61 | 0 | 25 |
| Total: | 327 | 2637 | 6230 | 60 | 527 |

Next, we analyzed the overlap of hybridization-dependent splicing events of ATXN3-AON in the three different cell lines, looking specifically at the 19 potential hybridization-dependent events that showed a larger absolute dPSI than the intended event in *ATXN3* in C1.1. From these 19 events, 5 were confirmed in cell lines C1.2 and C2 (Table 10). These events were located in *GOLGA3*, *FLOT1* and *RFX4*. These genes are involved in intracellular transport, vesicle-mediated transport and transcriptional regulation. None of these 5 events were found in the experiments with ATXN3-AON2, supporting our hypothesis that these are off-target events specific for ATXN3-AON.

**Table 10.** From the 19 potentially hybridization-dependent events with larger dPSI than the intended event of ATXN3-AON in C1.1, 5 were found in C1.2 and C2 as well. dPSI: delta percent spliced in.

| Gene | Exon coordinate | Flanking region | Event | dPSI C1.1 | dPSI C1.2 | dPSI C2 |
| --- | --- | --- | --- | --- | --- | --- |
| GOLGA3 | 12:132776634-132776756 | 12:132774496-132775305, 12:132777665-132777805 | Exon 21, between 22 (extended or last exon T203) and 19 | 0.73 | 0.32 | 0.66 |
| GOLGA3 | 12:132776958-132777090 | 12:132774496-132775305, 12:132777665-132777805 | Exon 20, between 22 (extended or last exon T203) and 19 | 0.60 | 0.23 | 0.50 |
| GOLGA3 | 12:132776634-132776756 | 12:132774496-132775305, 12:132776957-132777090 | Exon 21, between 22 (extended or last exon T203) and 20 | 0.52 | 0.24 | 0.45 |
| FLOT1 | 6:30730022-30730186 | 6:30727708-30728145, 6:30730427-30730565 | Exon 12 (from 13 exons), between exon 11 and 13 | 0.44 | 0.21 | 0.38 |
| RFX4 | 12:106681993-106682054 | 12:106654227-106654351, 12:106686883-106687097 | Exon 5 (from 18 exons), between exon 4 and 6) | 0.36 | 0.23 | 0.64 |

Finally, we investigated if there was any overlap in the 198 shared significant splicing events for ATXN3-AON with the 43 events that were identified for ATXN3-AON2 in the two cell lines C1.2 and C2. In principle, these overlapping splicing events could not be hybridization-dependent since the sequence of ATXN3-AON and ATXN3-AON2 are different while resulting in the same skip of ATXN3 exon 10. Only 6 splicing events were significantly induced by both ATXN3 targeting AONs (Fig 6B, Table S4), of which four of these events (in *NUCB1*, *SMARCC2*, *SGSM2* and *VEZF1*) were specific for the ATXN3-targeting AONs while the other 2 events could also be detected with the HTT-AON. All four ATXN3-related events showed a larger mean dPSI than the intended event in *ATXN3* in C1.1. These results illustrate that it is important to include proper control AONs to be able to determine which splicing events are AON-specific.

## Discussion

Splice-switching AONs are promising drugs for restoring protein expression or modifying the eventual protein to restore its function or reduce its toxicity (29). Given the current lack of *in silico* methods that adequately predict off-target splicing events, assessment of off-target effects of AONs in human cells using RNAseq is a promising approach (11). In this study, we assessed the validity of short-read RNAseq to identify transcriptome-wide off-target effects of three different splice-switching AONs targeting three different human transcripts and studied the role of hybridization-dependent off-target splicing events.

We observed substantial gene expression changes upon AON transfection, especially for the AON targeting *ATXN3*. Since the comparisons were made to a scrambled MOE-PS AON, general class effects of MOE-PS AONs and transfection-related changes in gene expression should have been cancelled out. Pathway analysis of differentially expressed genes that were shared by all three splice switching AONs, each with a different target RNA, affected pathways involved in RNA-related processes and neuronal function. These findings are in line with the study by Ottesen *et al*, where upon transfection of the *SMN2*-targeting 2’OMe-PS AON processes involved in transcription, RNA transport, cell signalling and synaptic activity were changed (30).

The efficiency of the intended splicing events in the target genes *APP*, *ATXN3* and *HTT* that we identified using splice analysis tools corresponded well with RT-PCR analysis. The overlap of splicing events identified by two different splice analysis tools was limited, as described before (15, 16). However, the effect sizes described by the two different tools correlated well. When we looked for potentially hybridization-dependent off-target effects, only a minority of events could be related to a predicted binding site. These splicing events were mainly AON-specific, supporting the hybridization-dependency of these events. In general, these potentially hybridization-dependent events showed a smaller effect size than the intended event, probably due to a decrease in exon skip efficiency with an increasing number of mismatches of the AONs to the RNA (11).

However, the majority of the splicing events that showed a larger effect size than the intended events could not be related to a predicted binding site, demonstrating that off-target splicing events are difficult to predict based on sequence homology. Furthermore, there was a large overlap between the larger events of the three AONs. As the comparison was made using the same scrambled AON as reference, these shared events might be induced by the scrambled AON. However, none of the shared events could be related to a predicted binding site of the scrambled AON. An alternative explanation could be that these shared events might be related to splice factors being directed away from their canonical splicing in the presence of a targeting AON or due to downstream effects of the intended events as APP, ATXN3 and HTT are involved in overlapping pathways.

The ATXN3-AON showed signs of toxicity as substantial cell death was observed after transfection and about 1.5 to 4 times as much DEGs and about 7 to 12 times as much splicing events as the APP-AON and the HTT-AON were detected. In addition, about 12% of all events detected upon ATXN3-AON transfection showed a larger effect size than the intended event in *ATXN3*. We investigated whether the AON-induced effects on gene expression and differential splicing were reproducible in multiple cell lines and with different sequencing depth and sample sizes between experiments.

About 3% of the splicing events identified in our initial experiment was reproduced in two other cell lines. Almost half of the validated splicing events showed a larger effect size than the intended effect in the first experiment, confirming that larger splicing events are more robust (11). The lack of detection of 97% of the events identified in the initial experiment is probably related to the lower sample size and sequencing depth in the validation experiment. When considering the splicing events identified in the cell line with the lowest sample size, 61% of the splicing could be validated in both other cell lines, suggesting that the events that are found with lower power are robust.

Regardless of the sample size and sequencing depth, comparing multiple AONs within an RNA sequencing experiment can provide useful information about AON toxicity. Similar to what was found in the first experiment, the effects for ATXN3-AON on gene expression and splicing were 5 times larger than for ATXN3-AON2, an AON that induces the same exon 10 skip but has a different sequence, and little overlap was found, confirming the toxic profile of ATXN3-AON. Since ATXN3-AON showed the highest number of predicted off-target binding sites this could indicate that the unfavourable effects of ATXN3-AON might be sequence-dependent. Moreover, among the splicing events that were consistently found in the different cell lines were 16 genes involved in the regulation of RNA splicing. Splicing events in these genes might alter their function, subsequently inducing other splicing events.

Taking into account the results from the current study, some improvements in the AON design and experimental set up to study off-target effects can be made. All four targeting AONs used in this study were designed according to the same guidelines (24) and only BLAST was used to check sequence specificity. Inclusion of predicted binding sites using GGGenome might give an more accurate indication of off-target binding sites. However, in line with previous studies, our results also showed that only a few of the GGGenome predicted binding sites of the AON actually induced a splicing event, demonstrating the difficulty to predict off-target effects based on sequence homology alone (8, 11). Furthermore, our study revealed several challenges of analysing off-target effects using short RNAseq, complicating the reproducibility and interpretation that require improvements in the experimental set-up. First of all, experimental factors such as cell culture and transfection protocols, time points, sequence of the scrambled AON, but also factors affecting the statistical analysis, such as sample size, RNAseq read depth and the settings in the splice analysis tools all affect the differentially expressed genes and significant splicing events detected. Secondly, the interpretation of the results is very difficult. Both on-target and off-target splicing events can lead to downstream transcriptional changes and splicing events. It is important to use a different AON with the same target event to be able to identify these on-target downstream effects. Furthermore, the use of multiple time points could also provide more insight in direct and indirect effects of AON transfection. Finally, it has been questioned how relevant *in vitro* transfection experiments are in the prediction of *in vivo* off-target effects as off-target effects were reduced when AONs were administered via free uptake instead of via transfection (8). Since free uptake is generally more predictive for *in vivo* uptake, it is recommended to include this as well (31).

Further investigation is required to determine the biological relevance of the off-target effects found in this study. Here, we studied the effects at 72 hours after transfection and the inclusion of earlier time points could shed light on direct and indirect effects of AON transfection. Also inclusion of cycloheximide, which inhibits RNA translation, could provide insight in induced splicing events that result in non-sense mediated decay as this process is translation-dependent {Pereverzev, 2015 #64}. As our study was limited to female cell lines, the effects on genes on the Y-chromosome should still be studied. Furthermore, the sequencing depth and sample size were lower in the validation experiment, reducing the power and sensitivity of the differential splicing analysis. Especially splicing events in genes with lower expressions could have been missed. Additionally, in order to link predicted binding sites to splicing events, we assumed that AON binding only has an effect on splicing when binding within the exon or surrounding introns and exons. We also only assessed binding sites with up to 3 mismatches with the AON sequences as these are most likely to bind, but even up to 5 mismatches seemed to result in splicing effects (11). Hence, we might not have covered all hybridization-dependent splicing effects.

Overall, our study illustrates the use of short-read RNAseq to identify and prioritize potential off-target effects of splice-switching AONs for further investigation of their biological relevance. Although the interpretation of short-read RNAseq is difficult, we and others showed that predicting hybridization-dependent off-target effects based on sequence homology is even more limited (8, 11). From the differential splicing events identified, only a minority could be explained by hybridization. An RNAseq experiment first of all allows the comparison of toxicity between different AONs within an experiment. From the three AONs studied, we identified one AON inducing many transcriptomic changes and splicing events compared to the two other AONs assessed, suggesting a toxic profile that corresponded with signs of cell death that was confirmed in a second experiment with two different cell lines. Secondly, we showed that RNA sequencing can be used to identify AON-specific off-target profiles, as the main splicing events that we prioritized could be validated with RT-PCR. Further investigation of these prioritized potential off-target effects would contribute to the development of safe and effective AONs.

## Acknowledgements

We would like to thank David van der Meer from GenomeScan for providing the information about the sequencing methods and the Dutch SCA1 giving circle for their contribution to the generation of the control iPSC cell lines.

## Author contributions

**ECK**: conceptualization, methodology, validation, formal analysis, investigation, writing – original draft, visualization, supervision, project administration, data curation. **LvdG**: investigation. **BAP**: investigation. **MRG**: investigation. **SK**: investigation. **LJAT**: conceptualization, methodology, investigation. **DC**: investigation. **RAMB**: conceptualization, writing – review & editing. **EM:** conceptualization, supervision, methodology. **HM**: conceptualization, methodology, investigation. **WMCvRM**: conceptualization, methodology, supervision, writing – review & editing, funding acquisition. All authors have read and approved the manuscript.

## Supplementary figures

**Figure S1.**
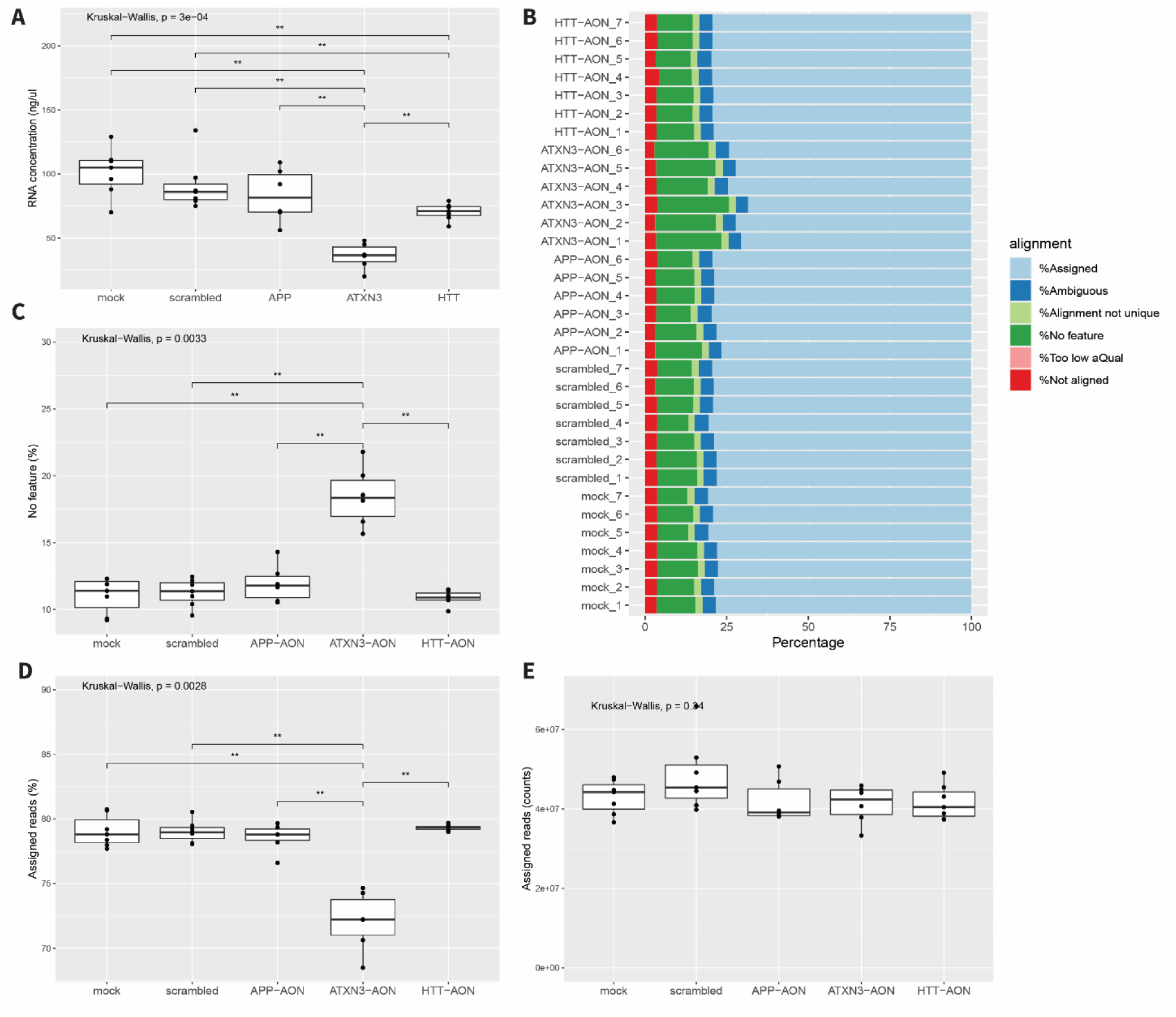
Results of RNA isolation and RNA sequencing. A) The RNA concentration upon isolation showed that especially transfection with the ATXN3-AON resulted in a lower RNA yield. B-D) Percentage-wise, more reads could not be assigned to a feature (C) and less reads were assigned to a gene (D) for ATXN3-AON-treated cells compared to the other conditions. E) Assigned reads per condition did not differ significantly between groups and showed an overall average of 43 million assigned reads. ** P < 0.01

**Figure S2.**
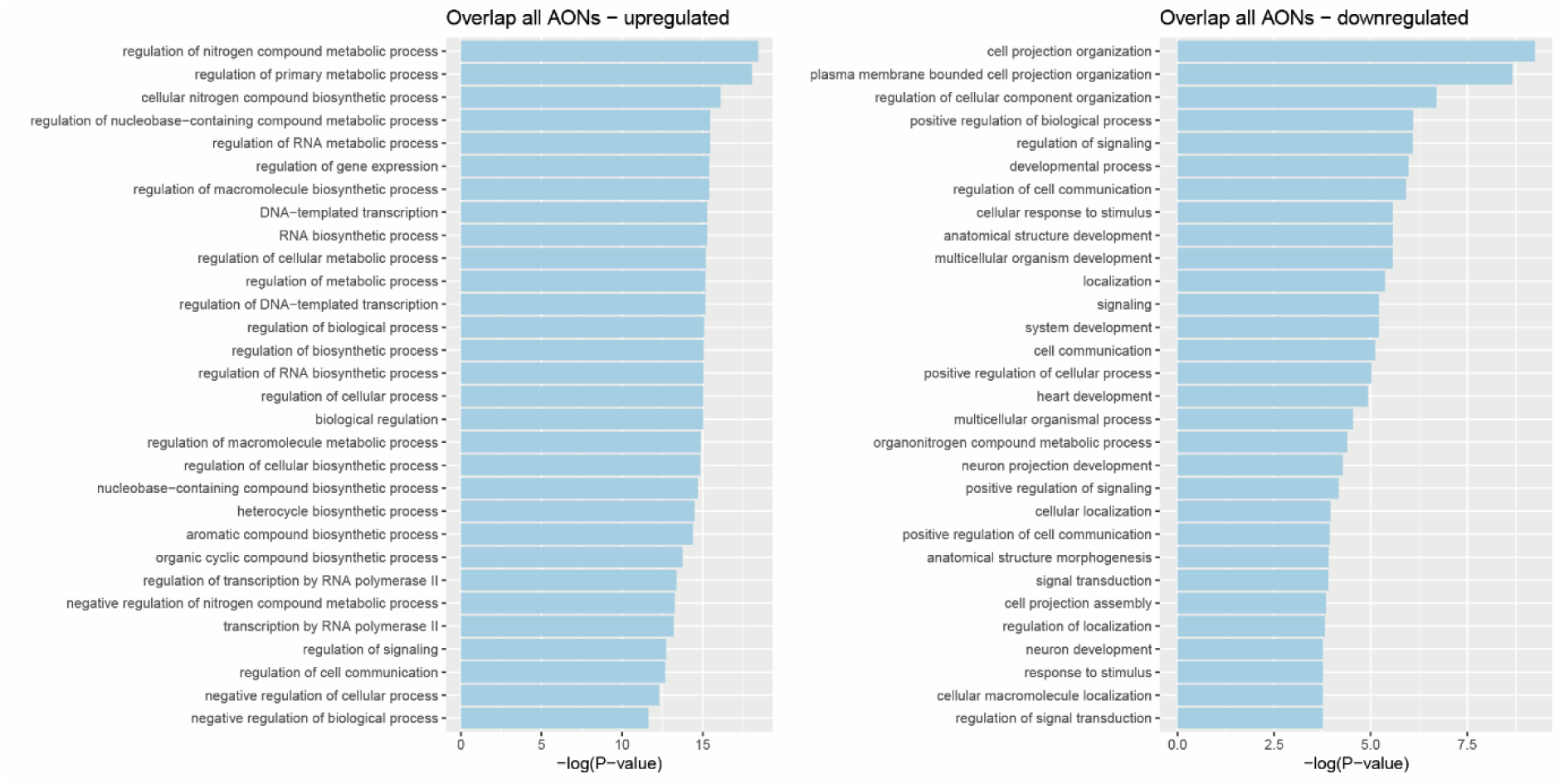
Top 30 enriched pathways for the up– and downregulated genes shared by all three targeting AONs.

**Figure S3.**
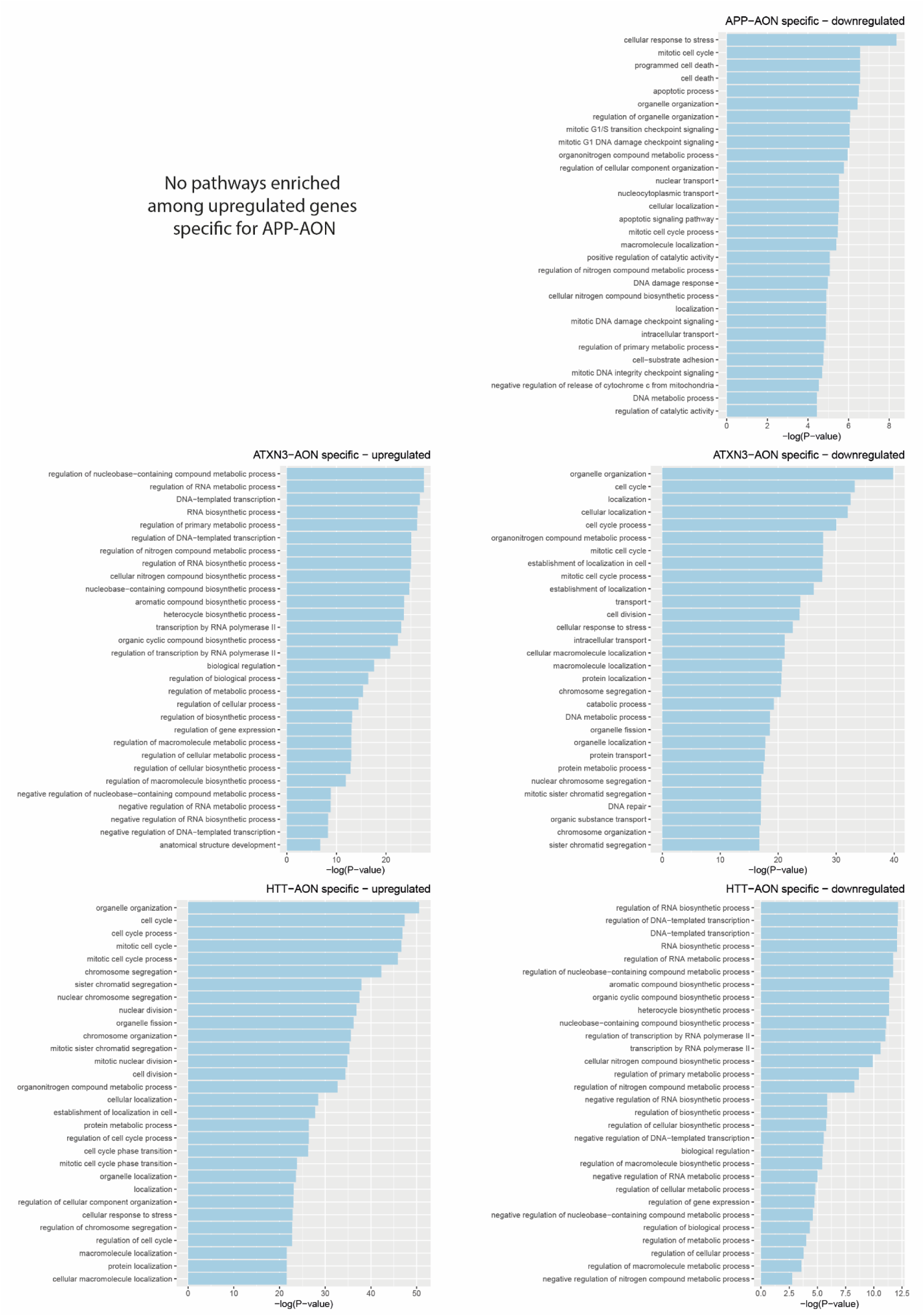
Top 30 enriched pathways for the up– and downregulated genes specific for each targeting AON.

**Figure S4.**
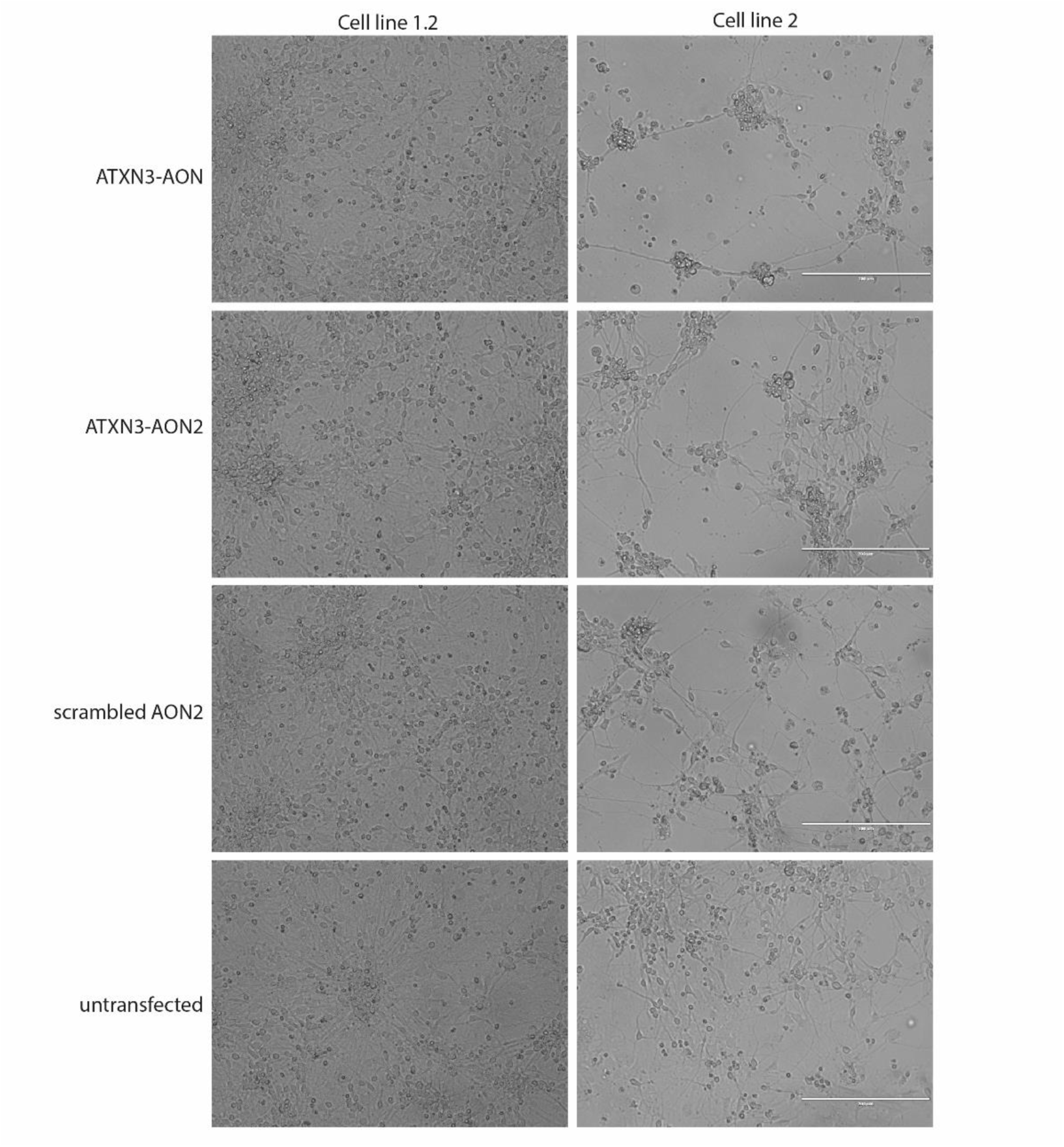
Pictures of iPSC-derived neurons 3 days after transfection. Especially transfection of ATXN3-AON led to cell death in cell line 2. Scale bar 200 µm; 20x magnification.

**Figure S5.**
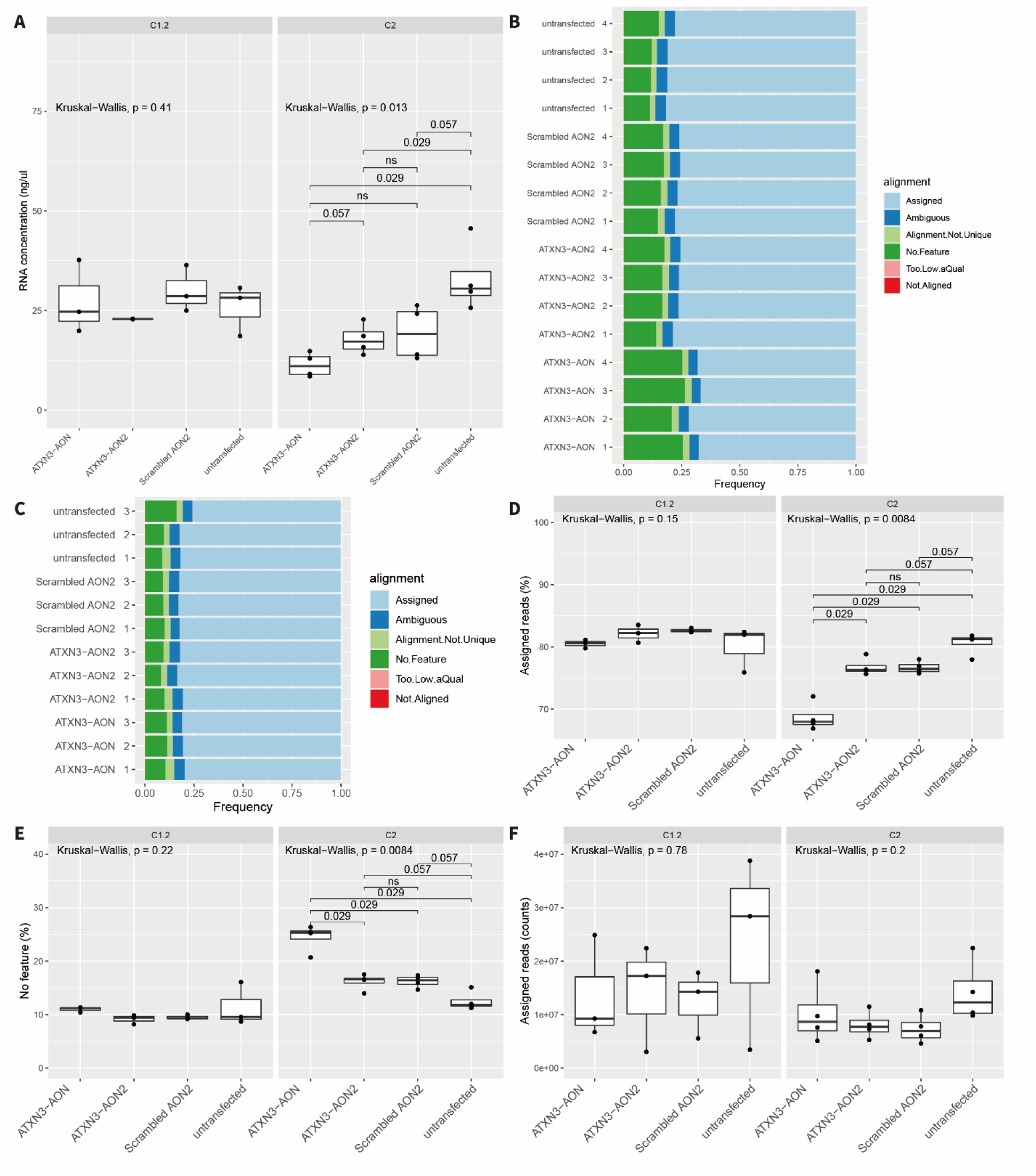
Results of RNA isolation and RNA sequencing in cell lines C1.2 and C2. A) Although the RNA concentration did not differ among the conditions in C1.2, especially transfection with the ATXN3-AON resulted in a lower RNA yield in C2. B) The alignment of reads showed less reads assigned to a gene and more reads showing no feature for ATXN3-AON-treated cells in C2 percentage-wise. C) This pattern was less obvious in C1.2. D-E) Consistently, statistical analysis of the percentage of reads assigned (D) and reads showing no feature (E) showed no difference among groups in C1.2, but a significantly lower percentage of reads assigned and percentage of reads showing no feature in C2. F) The number of assigned reads did not differ significantly among the groups in both cell lines.

**Figure S6.**
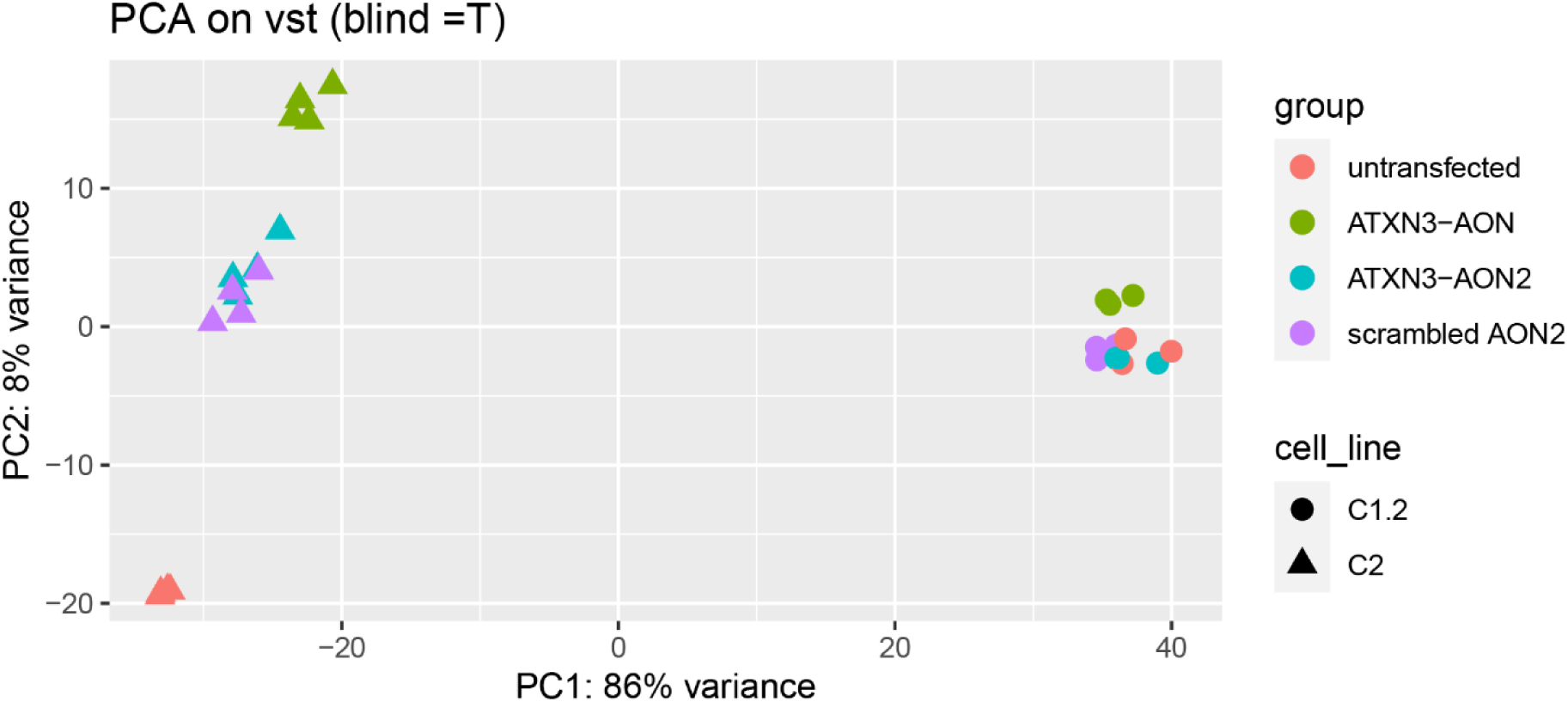
Principle component analysis of the samples included in the validation experiment in cell lines C1.2 and C2.

**Figure S7.**
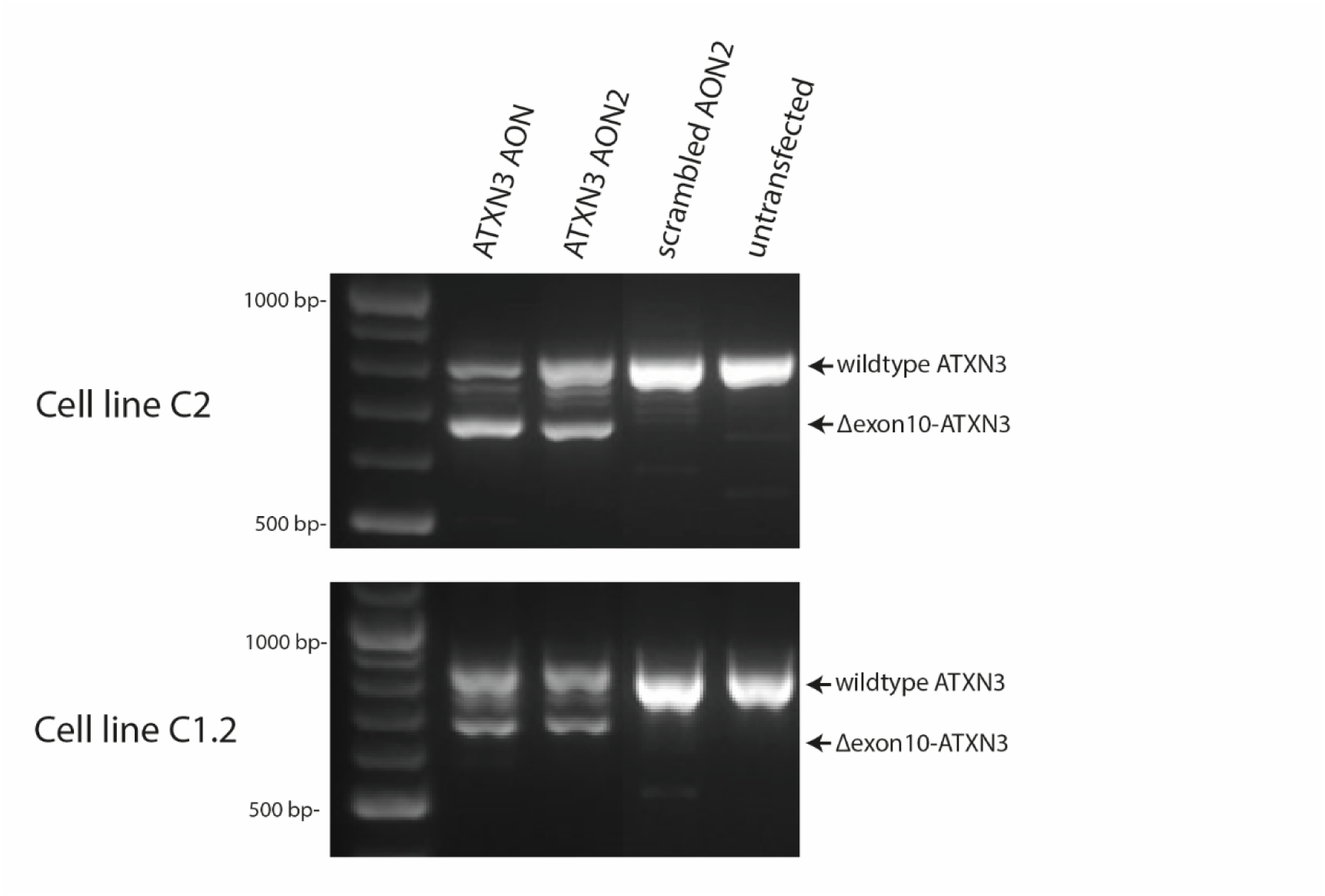
The intended event in ATXN3 was confirmed by RT-PCR for both ATXN3-targeting AONs in both cell lines.

**Figure S8.**
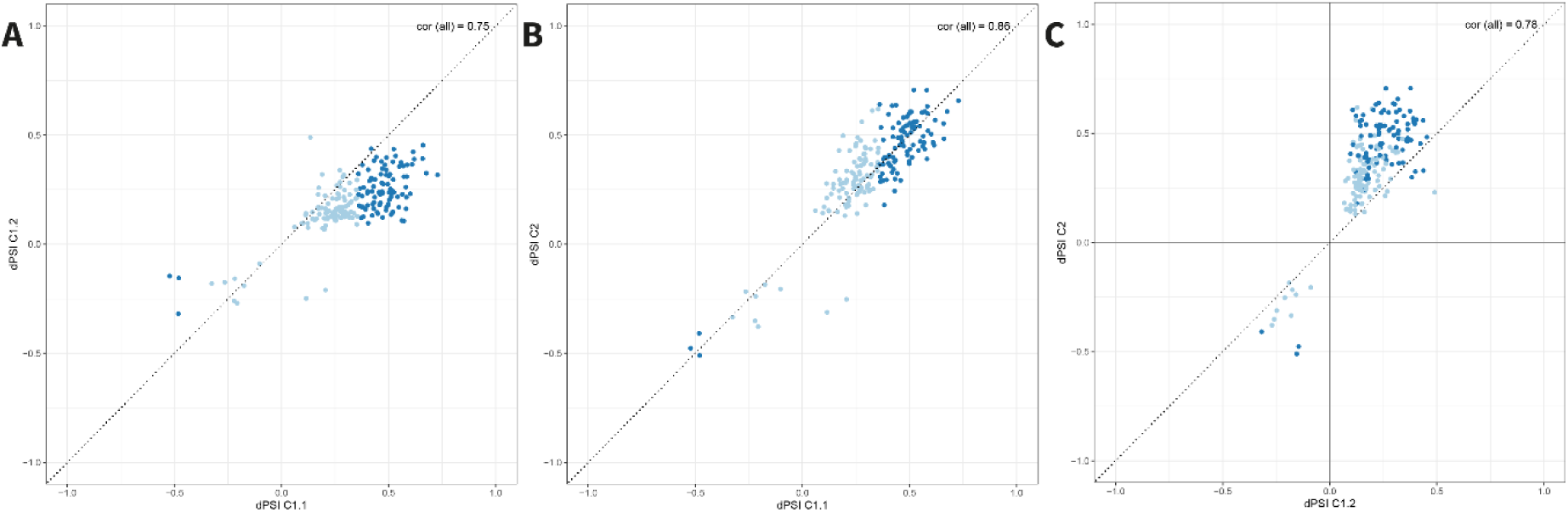
Correlation of the shared differential splicing events induced by ATXN3-AON transfection when compared to scrambled AON (C1.1) or scrambled AON2 (C1.2 and C2). dPSI: delta percent spliced in. AON-specific events larger than the intended event in *ATXN3* identified in C1.1 are highlighted in darkblue. Correlations (cor) were determined using all shared events.

## Supplementary tables

**Table S1.** The significant events identified with rMATS and Whippet in the target genes represented the designed exon skip events. SE: exon skipping; A5SS: alternative donor splice site; FDR: false discovery rate; PSI: percent spliced in; dPSI: delta PSI; Scr: scrambled AON; Prob: probability.

| Scrambled AON vs. APP-AON |  |  |  | rMATS |  |  |  | Whippet |  |  |  | Average |
| --- | --- | --- | --- | --- | --- | --- | --- | --- | --- | --- | --- | --- |
| Gene | Type | Flanks | Coordinates | FDR | PSI Scr | PSI AON | dPSI | Prob | PSI Scr | PSI AON | dPSI | dPSI |
| APP | SE | 21:25880550-25881771, 21:25897573-25897673 | 21:25891722-25891868 | 0 | 1.0 | 0.575 | 0.425 | 1 | 0.999 93 | 0.611 31 | 0.388 62 | 0.40681 |
| Scrambled AON vs. ATXN3-AON |  |  |  | rMATS |  |  |  | Whippet |  |  |  | Average |
| Gene | Type | Flanks | Coordinates | FDR | PSI Scr | PSI AON | dPSI | Prob | PSI Scr | PSI AON | dPSI | dPSI |
| ATXN3 | SE | 14:92058552-92064414, 14:92080965-92081061 | 14:92070935-92071053 | 0 | 1.0 | 0.605 | 0.395 | 0.979 | 0.981 89 | 0.661 58 | 0.320 31 | 0.357655 |
| Scrambled AON vs. HTT-AON |  |  |  | rMATS |  |  |  | Whippet |  |  |  | Average |
| Gene | Type | Flanks | Coordinates | FDR | PSI Scr | PSI AON | dPSI | Prob | PSI Scr | PSI AON | dPSI | dPSI |
| HTT | A5SS | 4:3129924-3130047 | 4:3127470-3127604 | 0 | 0.996 | 0.551 | 0.445 | 1 | 0.988 6 | 0.667 94 | 0.320 66 | 0.38283 |

**Table S2.**
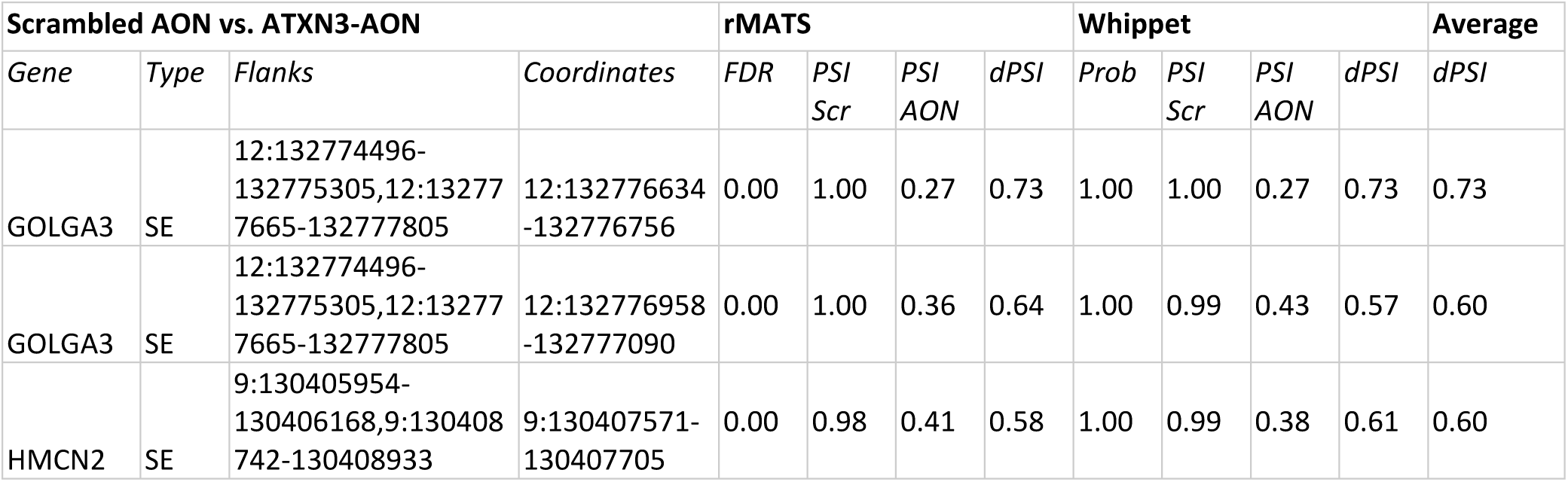
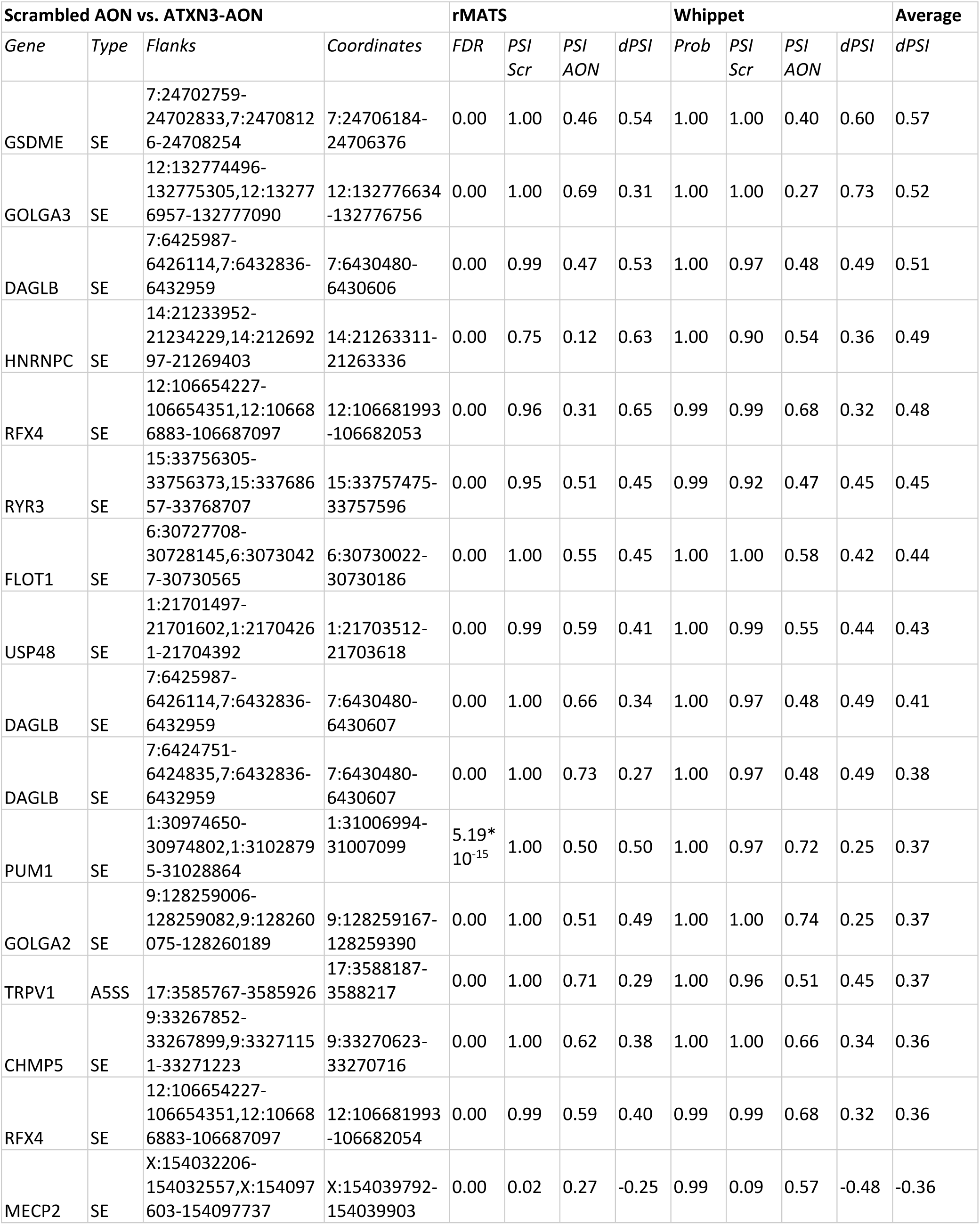
Potentially hybridization-dependent events induced by ATXN3-AON that showed a larger absolute dPSI than the intended event in ATXN3. SE: exon skipping; A5SS: alternative donor splice site; FDR: false discovery rate; PSI: percent spliced in; dPSI: delta PSI; Scr: scrambled AON; Prob: probability.

**Table S3.** Shared non-hybridization-dependent events larger than the intended events of the AONs. SE: exon skipping; dPSI: delta PSI.

|  |  |  |  | APP-AON | ATXN3-AON | HTT-AON |
| --- | --- | --- | --- | --- | --- | --- |
| Gene | Type | Flanks | Coordinates | dPSI | dPSI | dPSI |
| ARHGAP29 | SE | 1:94189925-94190083,1:94202543-94202732 | 1:94201720-94201857 | -0.442 | -0.619 | -0.442 |
| CDC16 | SE | 13:114236798-114236896,13:114239349-114239490 | 13:114238990-114239028 | -0.489 | -0.483 | -0.489 |
| CNN3 | SE | 1:94901668-94901785,1:94903121-94903188 | 1:94902121-94902258 | -0.533 | -0.532 | -0.533 |
| IFTAP | SE | 11:36610080-36610239,11:36648015-36650464 | 11:36633284-36633438 | -0.442 | -0.388 | -0.442 |
| INTS6L | SE | X:135575083-135575226,X:135579787-135580162 | X:135577193-135577295 | -0.525 | -0.520 | -0.525 |
| LGR4 | SE | 11:27380639-27380711,11:27382187-27382256 | 11:27380895-27380966 | -0.425 | -0.425 | -0.425 |
| PIKFYVE | SE | 2:208350770-208350947,2:208352653-208352782 | 2:208351352-208351455 | -0.453 | -0.466 | -0.453 |
| PRXL2C | SE | 9:96651389-96651495,9:96654704-96654773 | 9:96651659-96651712 | -0.434 | -0.432 | -0.434 |
| RAPGEF2 | SE | 4:159186641-159186712,4:159210499-159210583 | 4:159193200-159193256 | -0.586 | -0.584 | -0.586 |
| VTI1B | SE | 14:67659730-67659922,14:67674374-67674632 | 14:67662477-67662535 | -0.582 | -0.513 | -0.582 |
| YME1L1 | SE | 10:27136275-27136385,10:27145427-27145590 | 10:27142387-27142485 | -0.556 | -0.457 | -0.556 |
| YY1AP1 | SE | 1:155679408-155679512,1:155688070-155688201 | 1:155680416-155680456 | -0.541 | -0.508 | -0.541 |

**Table S4.** Shared splicing events by ATXN3-AON and ATXN3-AON2. SE: exon skipping; dPSI: delta PSI.

| Gene | Type | Flanks | Coordinates | dPSI C1.1 | dPSI C1.2 | dPSI C2 |
| --- | --- | --- | --- | --- | --- | --- |
| BCLAF1 | SE | 6:136261264-136261477,6:136268161-136268339 | 6:136267029-136267175 | 0.205 | -0.210 | -0.252 |
| BECN1 | SE | 17:42814523-42814673,17:42818220-42818415 | 17:42815908-42816054 | 0.237 | 0.210 | 0.324 |

| *Gene* | *Type* | *Flanks* | *Coordinates* | **dPSI C1.1** | **dPSI C1.2** | **dPSI C2** |
| --- | --- | --- | --- | --- | --- | --- |
| NUCB1 | SE | 19:48921153-48921324,19:48922317-48923255 | 19:48921869-48921932 | 0.522 | 0.296 | 0.528 |
| SMARCC2 | SE | 12:56164302-56164731,12:56168059-56168194 | 12:56165318-56165699 | 0.468 | 0.281 | 0.519 |
| SGSM2 | SE | 17:2372952-2373081,17:2375491-2375875 | 17:2373331-2373513 | 0.376 | 0.220 | 0.365 |
| VEZF1 | SE | 17:57971551-57974900,17:57980602-57980786 | 17:57979152-57979313 | 0.582 | 0.162 | 0.390 |

