## Supplementary File 1 for "Determining off-target effects of splice-switching antisense oligonucleotides using short read RNAseq in neuronally differentiated human induced pluripotent stem cells"

### Supplementary File I: Optimization of differential splicing detection

We selected rMATS and Whippet as methods to detect differential splicing as rMATS performs well on detection of novel events and the combination of both methods has been shown to improve the FDR and a pipeline to compare the methods is already established. During the setting optimization, the designed event that was validated with PCR was used as positive control and the aim was to focus on robust effects and reduce the noise as much as possible.

#### rMATS optimization

We started optimization based on the comparison between samples treated with scrambled AON and APP-AON, enabling --*variable-read-length* and *--allow-clipping*. Since we were interested in robust and biologically relevant splicing alterations, we followed the recommendation from Liang et al to change the cutoff splicing difference (*cstat*) value to 5% instead of using the default (0.01%). The change of this setting drastically decreased the number of significant effects (based on junction counts only) from 14957 events to 1540 events. As the AONs could induce the use of an unannotated splice site, we allowed *--novelSS*. (Indeed, the positive effect for the HTT-AON was only detected when novel splice sites were enabled.) When allowing novel splice sites, 3832 significant events were detected for APP-AON. These settings were applied to all three comparisons (Table 1, ‘setting 1’). With these settings, about 87% of all reads were used in the analysis. Next, we tested the effect of changing the library type to fr-firststrand, as the data was strand-specific, and setting the anchor length at 15 instead of 1 to reduce false positives. Also, we tried the use of a gtf-file without lowly expressed genes as recommended (1). The positive events were found with all setting variations. We combined the libType *fr-*firststrand, anchor length 15 and use of a filtered gtf-file into ‘setting 2’ (Table 1).

Table 1 Total and significant events found with use of different settings for the comparison of samples treated with scrambled AON and APP-AON (APP), ATXN3-AON (ATXN3) or HTT-AON (HTT).

| APP | Setting 1 | | Fr-firststrand | | Anchor length 15 | | Filtered GTF | | Setting 2 | |
| --- | --- | --- | --- | --- | --- | --- | --- | --- | --- | --- |
| Type | Total | Sign | Total | Sign | Total | Sign | Total | Sign | Total | Sign |
| SE | 306087 | **2476** | 304885 | **2468** | 291634 | **2225** | 303695 | **2460** | 288297 | **2206** |
| A5SS | 94425 | **281** | 93943 | **272** | 86879 | **195** | 93325 | **281** | 85419 | **190** |
| A3SS | 140043 | **344** | 139452 | **336** | 127348 | **230** | 138821 | **340** | 125682 | **227** |
| MXE | 113743 | **624** | 112914 | **614** | 110034 | **565** | 111226 | **589** | 106832 | **550** |
| RI | 41946 | **107** | 41749 | **107** | 39474 | **122** | 41554 | **106** | 38876 | **119** |
| ATXN3 | Setting 1 | | Fr-firststrand | | Anchor length 15 | | Filtered GTF | | Setting 2 | |
| Type | Total | Sign | Total | Sign | Total | Sign | Total | Sign | Total | Sign |
| SE | 395229 | **24742** | 393665 | **24640** | 377224 | **22352** | 392383 | **24614** | 373119 | **22096** |
| A5SS | 110667 | **3409** | 110123 | **3318** | 101570 | **2685** | 109516 | **3350** | 99980 | **2543** |
| A3SS | 165365 | **3769** | 164627 | **3688** | 149712 | **2637** | 164035 | **3745** | 147797 | **2507** |
| MXE | 175568 | **9467** | 174312 | **9401** | 170113 | **8352** | 171819 | **9194** | 165286 | **8057** |
| RI | 48434 | **3016** | 48185 | **2749** | 45689 | **2706** | 48008 | **3005** | 44994 | **2417** |
| HTT | Setting 1 | | Fr-firststrand | | Anchor length 15 | | Filtered GTF | | Setting 2 | |
| Type | Total | Sign | Total | Sign | Total | Sign | Total | Sign | Total | Sign |
| SE | 338986 | **4232** | 337540 | **4196** | 322618 | **3707** | 336172 | **4196** | 318619 | **3644** |
| A5SS | 99635 | **501** | 99192 | **495** | 91706 | **375** | 98507 | **490** | 90223 | **351** |
| A3SS | 150554 | **587** | 149870 | **578** | 136884 | **399** | 149298 | **585** | 135066 | **385** |
| MXE | 132595 | **1100** | 131739 | **1090** | 128597 | **943** | 129479 | **1070** | 124778 | **913** |
| RI | 43629 | **247** | 43436 | **208** | 41058 | **208** | 43224 | **243** | 40449 | **183** |

Next, we compared the detection of the positive events between setting 1 and 2 after filtering the rMATS results on informative reads present for each event (at least 10 reads for each sample, Figure 1). Whereas the detection of the positive APP- and HTT-AON effect seemed to be similar, the detection of the ATXN3-AON event was more specific.

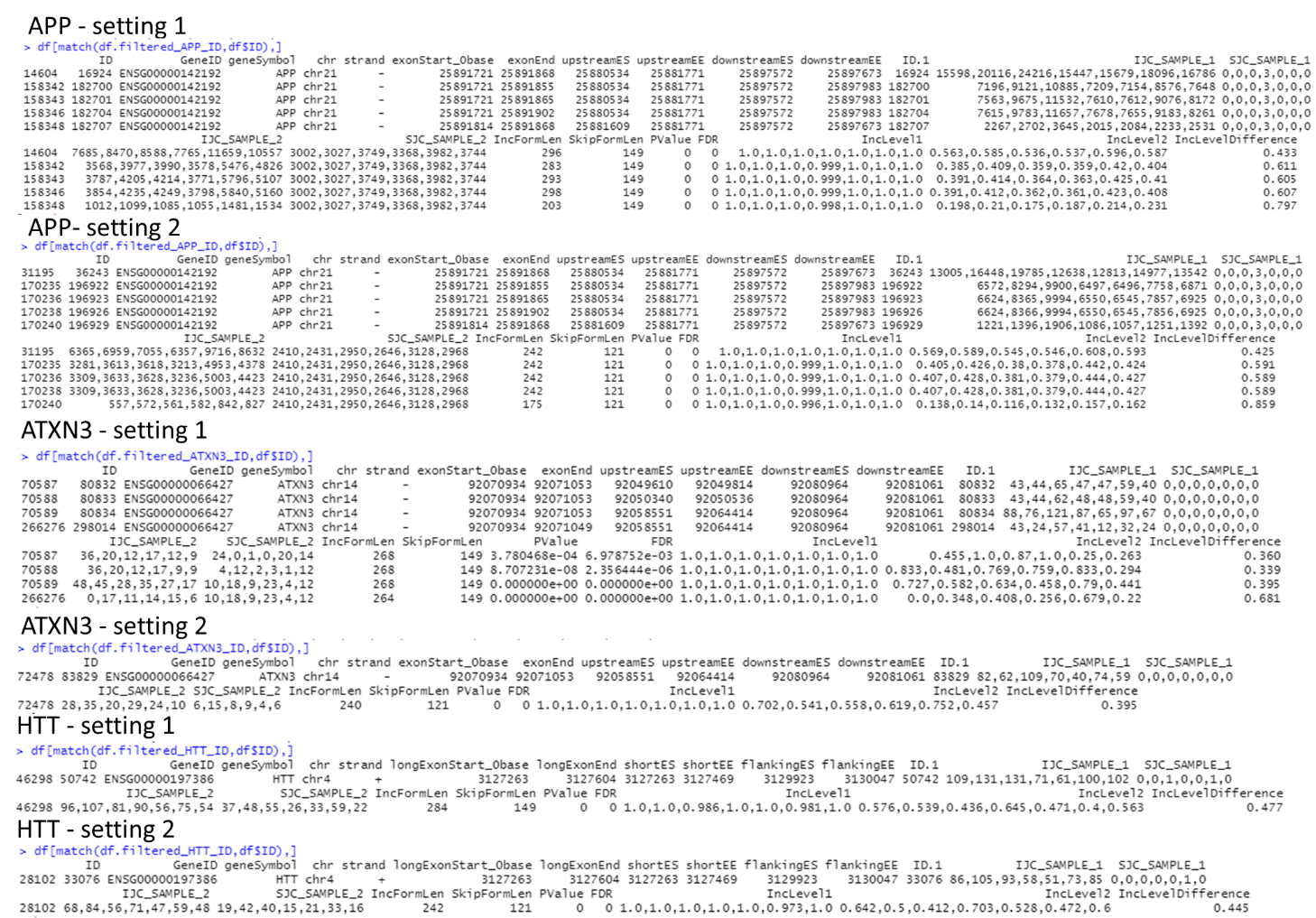

Figure 1 Detection of positive events using setting 1 or 2.

#### Whippet optimization

The pipeline of Whippet consists of three scripts: making an index, quantifying each sample and calculating the delta PSI between two sample groups. The optimization of Whippet was aimed at getting results comparable to the results of rMATS while adjusting the settings to the tool’s own needs to get robust results.

To allow detection of novel events, we supplied a merged bam-file containing the samples treated with scrambled AON and a targeting AON (APP-, ATXN3- or HTT-AON) to include de novo splice sites in the index. The setting *–suppress-low-tsl* was enabled to reduce noise. Two settings could be adjusted influencing the detection of novel splice sites: the minimum number of reads supporting a splice site from BAM to be included in the index (*--bam-min-reads*) and whether splicing reads with two de novo splice sites were allowed (*--bam-both-novel*). As expected, the number of de novo splice sites was higher when *--bam-both-novel* was enabled and decreased with the use of a higher minimum reads (Table 2). For all options tried, the novel splice site introduced by HTT-AON could be detected.

Table 2 Detection of splice-sites in the GTF file and based on the BAM-file using different settings.

| **Bam-min-reads** | **Bam-both-novel** | **+** | **-** |
| --- | --- | --- | --- |
| **2** | Scrapp | 187150 BAM  533929 GTF | 78624 BAM  533929 GTF |
| **2** | Scrhtt | 190956 BAM  533929 GTF | 80555 BAM  533929 GTF |
| **3** | Scrapp | 124833 BAM  533929 GTF | 52742 BAM  533929 GTF |
| **6** | Scrapp | 72367 BAM  533929 GTF | 26848 BAM  533929 GTF |
| **6** | Scrhtt | 73502 BAM  533929 GTF | 27571 BAM  533929 GTF |
| **6** | Scratxn3 | 85369 BAM  533929 GTF | 34813 BAM  533929 GTF |

For the final index, we enabled *--bam-both-novel* to include all potential AON-related effects and we chose a minimum reads of 6, since each group had at least 6 samples. Also, we used the same filtered GTF-file as used for setting 2 of rMATS.

However, with these settings, the positive event for ATXN3-AON could not be found. The only event found for exon 10 contained only a part of the exon and was classified as tandem alternative polyadenylation site (use of exon 10 as last exon instead of exon 11), which is not the case, with a low probability and very small deltaPSI. Also with adjusting the minimum reads threshold in the script to calculate the deltaPSI from default (5) to 1, since the expression of ATXN3 was low, did not change the outcome. The first nucleotides of exon 10 (92070935:92071037) were not quantified by default (node without a simple type, NA) which could contribute to the failure to detect the exon skip. As the only remedy to change this is simplifying the input reference annotation, we looked at the number of different isoforms annotated for ATXN3. Compared to APP (20 isoforms) and HTT (23 isoforms), the number of ATXN3 isoforms (54) is indeed quite high. The high number of isoforms might especially be complicated as Whippet distinguishes a larger number of event types compared to rMATS. In combination with the low expression of ATXN3, this might complicate proper splice detection. Therefore, we followed the advice to use a simplified reference annotation and tried the use of isoforms annotated with gencode-basic only. Now, the skip of exon 10 was finally detected. To be consistent, we used the gencode-basic gtf for all three comparisons.

#### Assessing overlap

To see whether the use of a gtf subset (gencode-basic) improved the performance of Whippet in particular or whether a consistent use of the same gtf as input for both Whippet and rMATS was preferable, we compared the overlap of the significant results for both tools with different input gtf-files (Table 3). In the overlap, only the event types shared by both tools are present: SE, A5SS, A3SS and RI. The overlap of significant events is only about 25%, which is consistent with literature (2). The use of a basic gtf-file in Whippet increases the overlap with rMATS. For rMATS, the use of a basic gtf-file does not change much in the overlap. Especially considering the larger events with an absolute deltaPSI >0.3 in both tools, the overlap is the same. This is not surprising as splice sites that have been removed from the gtf-file can still be detected as novel splice site as de novo site detection is enabled. To focus on robust findings, we chose to use the basic gtf-file for rMATS as well.

Table 3 Overlap of significant events found by rMATS or Whippet while using the gtf-file filtered on expression levels or also filtered for genes annotated as gencode-basic ('basic gtf'). The significant events obtained for the different comparisons are listed in the first column for rMATS and the second row for Whippet (containing only the four event types occuring in both tools). The overlap presented is the 1) number of overlapping events, 2) percentage of the significant events found by the tools on their own and 3) the number of overlapping events with a deltaPSI >0.3 in both tools.

|  | ***Whippet*** | | | ***Whippet – basic*** | | |
| --- | --- | --- | --- | --- | --- | --- |
| ***rMATS*** | **APP: 3029** | **ATXN3: 20003** | **HTT: 4334** | **APP: 2703** | **ATXN3: 18703** | **HTT: 4107** |
| **APP: 1813** | 436  14-24%  67 |  |  | 467  17-26%  72 |  |  |
| **ATXN3: 22671** |  | 5911  26-30%  747 |  |  | 6088  27-33%  789 |  |
| **HTT: 3225** |  |  | 768  18-24%  55 |  |  | 839  20-26%  59 |
| ***rMATS – basic gtf*** |  | | | ***Whippet – basic*** | | |
| **APP: 1738** |  |  |  | 463  17-27%  73 |  |  |
| **ATXN3: 21300** |  |  |  |  | 6012  28-32%  788 |  |
| **HTT: 3118** |  |  |  |  |  | 843  21-27%  60 |

Finally, we wondered whether disabling the setting *–bam-both-novel* would improve the performance of Whippet as the tool’s authors have shown that this function increases the FDR and our main focus is on hybridization-dependent effects and the binding of an AON is more likely to induce one novel splice site. Hence, we run the script for Whippet again now without *–bam-both-novel*, both using the ‘normal’ gtf as the basic gtf, and compared the overlap with rMATS (Table 4). Although there were less significant events identified by Whippet when disabling *–bam-both-novel*, the overlap with the rMATS results was higher. Thus, we chose to disable this option while generating the index for Whippet. The numbers of novel splice sites detected in the BAM files with or without *–bam-both-novel* for the normal gtf or basic gtf are listed in Table 5.

Table 4 Overlap of significant events found by rMATS or Whippet (disabled –bam-both-novel) while using the gtf-file filtered on expression levels or also filtered for genes annotated as gencode-basic ('basic gtf'). The significant events obtained for the different comparisons are listed in the first column for rMATS and the second row for Whippet (containing only the four event types occuring in both tools). The overlap presented is the 1) number of overlapping events, 2) percentage of the significant events found by the tools on their own and 3) the number of overlapping events with a deltaPSI >0.3 in both tools.

|  | ***Whippet – bbn off*** | | | ***Whippet – basic – bbn off*** | | |
| --- | --- | --- | --- | --- | --- | --- |
| ***rMATS*** | **APP: 2720** | **ATXN3: 18015** | **HTT: 3959** | **APP: 2445** | **ATXN3: 16745** | **HTT: 3712** |
| **APP: 1813** | 462  17-25%  69 |  |  | 491  20-27%  74 |  |  |
| **ATXN3: 22671** |  | 6136  27-34%  764 |  |  | 6280  28-38%  819 |  |
| **HTT: 3225** |  |  | 816  21-25%  63 |  |  | 860  23-27%  63 |
| ***rMATS – basic gtf*** |  | | | ***Whippet – basic*** | | |
| **APP: 1738** |  |  |  | 489  20-28%  75 |  |  |
| **ATXN3: 21300** |  |  |  |  | 6230  29-37%  820 |  |
| **HTT: 3118** |  |  |  |  |  | 867  23-28%  64 |

Table 5 Detection of splice-sites in the GTF file and based on the BAM-file using the normal or basic GTF file, with or without –bam-both-novel. All indexes are based on –bam-min-reads of 6.

| **GTF (filtered)** | **Bam-both-novel** | **+** | **-** |
| --- | --- | --- | --- |
| **Normal** | Scrapp | 61618 BAM  429970 GTF | 24243 BAM  429970 GTF |
| **Normal** | Scratxn3 | 73623 BAM  429970 GTF | 31791 BAM  429970 GTF |
| **Normal** | Scrhtt | 62678 BAM  429970 GTF | 24890 BAM  429970 GTF |
| **Basic** | Scrapp | 60291 BAM  384618 GTF | 28513 BAM  384618 GTF |
| **Basic** | Scratxn3 | 75158 BAM  384618 GTF | 36392 BAM  384618 GTF |
| **Basic** | Scrhtt | 61384 BAM  384618 GTF | 29181 BAM  384618 GTF |

#### Final settings

The final settings have been described in the method section. Table 6 lists the number of events obtained using the basic gtf-file for both rMATS and Whippet for all event types detected by the tools, whereas the overlap of the event types assessed by both tools is listed in Table 7.

Table 6 Final number of (significant) results obtained by rMATS (after filtering results on read count) and Whippet per event type.

| rMATS | APP | | ATXN3 | | HTT | |
| --- | --- | --- | --- | --- | --- | --- |
| Type | Total | Sign | Total | Sign | Total | Sign |
| SE | 184479 | **1527** | 229819 | **17347** | 200313 | **2746** |
| A5SS | 50738 | **88** | 56311 | **1503** | 52353 | **145** |
| A3SS | 76760 | **80** | 85336 | **1320** | 80983 | **165** |
| MXE | 69435 | **414** | 103745 | **6902** | 82235 | **636** |
| RI | 17039 | **43** | 18880 | **1130** | 17329 | **62** |
| Whippet | APP | | ATXN3 | | HTT | |
| Type | Total | Sign | Total | Sign | Total | Sign |
| AA | 8579 | **210** | 9205 | **1152** | 8535 | **223** |
| AD | 16126 | **350** | 19842 | **3462** | 16682 | **375** |
| AF | 3005 | **32** | 2943 | **222** | 3003 | **71** |
| AL | 1035 | **23** | 951 | **75** | 1005 | **33** |
| CE | 146922 | **1822** | 148228 | **11629** | 146730 | **3018** |
| RI | 2715 | **63** | 2838 | **502** | 2717 | **96** |
| TE | 27079 | **515** | 27079 | **1256** | 26965 | **513** |
| TS | 20169 | **277** | 19852 | **454** | 20064 | **351** |

Table 7 Number of significant results obtained by rMATS and Whippet and overlap of significant results per event type. SE/CE: exon skipping; A5SS/AD: alternative splice donor site; A3SS/AA: alternative splice acceptor site; RI: retained intron.

|  | **APP-AON** | | | **ATXN3-AON** | | | **HTT-AON** | | |
| --- | --- | --- | --- | --- | --- | --- | --- | --- | --- |
| Type | *rMATS* | *Whippet* | ***Overlap*** | *rMATS* | *Whippet* | ***Overlap*** | *rMATS* | *Whippet* | ***Overlap*** |
| *SE/CE* | 1527 | 1822 | 468 | 17347 | 11629 | 5741 | 2746 | 3018 | 842 |
| *A5SS/AD* | 88 | 350 | 9 | 1503 | 3462 | 286 | 145 | 375 | 14 |
| *A3SS/AA* | 80 | 210 | 10 | 1320 | 1152 | 142 | 165 | 223 | 9 |
| *RI* | 43 | 63 | 2 | 1130 | 502 | 61 | 62 | 96 | 2 |
